# A genetic model of congenital intestinal atresia in medaka (*Oryzias latipes*) implicates Mypt1 in epithelial organisation

**DOI:** 10.1101/2021.12.10.472183

**Authors:** Daisuke Kobayashi, Akihiro Urasaki, Tetsuaki Kimura, Satoshi Ansai, Kazuhiko Matsuo, Hayato Yokoi, Shigeo Takashima, Tadao Kitagawa, Takahiro Kage, Takanori Narita, Tomoko Jindo, Masato Kinoshita, Kiyoshi Naruse, Yoshiro Nakajima, Masaki Shigeta, Shinichiro Sakaki, Satoshi Inoue, Rie Saba, Kei Yamada, Takahiko Yokoyama, Yuji Ishikawa, Kazuo Araki, Yumiko Saga, Hiroyuki Takeda, Kenta Yashiro

**Author notes:** Address correspondence to: Daisuke Kobayashi, Hiroyuki, Takeda and Kenta Yashiro. Graduate School of Science, Technology and Innovation, Kobe University (Akihiro Urasaki). Faculty of Life Sciences, Kyoto Sangyo University, Kyoto 603-8555, Japan (Hiroyuki Takeda).

## Abstract

Congenital intestinal atresia (IA) is a birth defect characterised by the absence or closure of part of the intestine. Although genetic factors are implicated, mechanistic understanding has been hindered by the lack of suitable animal models. Here, we describe a medaka (*Oryzias latipes*) mutant, generated by N-ethyl-N-nitrosourea (ENU) mutagenesis, that develops IA during embryogenesis. Positional cloning identified a nonsense mutation in *mypt1,* encoding *myosin phosphatase target subunit 1*. Mutant embryos exhibited ectopic accumulation of F-actin and phosphorylated myosin regulatory light chain (Mrlc) in the intestinal epithelium, consistent with disrupted actomyosin regulation. These cytoskeletal abnormalities were accompanied by epithelial disorganisation without notable alterations in cell proliferation, motility, or apoptosis. Inhibition of *myh11a*, encoding smooth muscle (SM) myosin heavy chain, ameliorated the IA phenotype but Blebbistatin treatment completely rescued the defect, suggesting a non-contractile role prior to SM maturation. Together, these findings demonstrate that *mypt1* loss disrupts intestinal morphogenesis through actomyosin dysregulation. Given the recent clinical identification of IA associated with MYPT1 mutations, this medaka model offers a valuable platform to investigate the developmental and molecular basis of *MYPT1*-associated IA in human.

## Introduction

Intestinal atresia (IA) is a congenital defect characterised by complete occlusion of the intestinal lumen, with an estimated incidence of 1.3 to 2.9 per 10,000 live births (Frischer and Azizkhan, 2012). Affected neonates require immediate surgical intervention to restore gastrointestinal continuity. Historically, IA has been attributed to in utero vascular accidents that impair blood supply to the developing gut (Louw, 1959; Louw and Barnard, 1955) . However, familial clustering and increased prevalence in individuals with trisomy 21 strongly suggest a genetic contribution to disease aetiology (Celli, 2014; Gupta et al., 2013; Louw, 1959; Shorter et al., 2006).

Mouse models support for this genetic hypothesis. Targeted deletion of *Fgfr2IIIb* or its ligand *Fgf10* results in IA in the absence of mesenteric vascular occlusion (Fairbanks et al., 2004a; Fairbanks et al., 2004b; Fairbanks et al., 2005; Kanard et al., 2005) . More recently, mutations in *PPP1R12A*, which encodes myosin phosphatase target subunit 1 (MYPT1), have been identified in human patients with IA and other developmental anomalies, including holoprosencephaly, urogenital malformations and persistent Müllerian duct syndrome (PMDS).(Hughes et al., 2020) (Picard et al., 2022). Despite these insights, mechanistic studies of MYPT1-related IA remain limited, as conventional *Mypt1* knockout mice exhibit early embryonic lethality (Okamoto et al., 2005). Actomyosin dynamics play a vital role in morphogenesis (Munjal and Lecuit, 2014). Non-muscle myosin II (NMII)-mediated contractility regulates cell shape, adhesion, migration, and tissue organisation (Agarwal and Zaidel-Bar, 2018; Munjal and Lecuit, 2014; Munjal et al., 2015; Vasquez et al., 2014). Phosphorylation of myosin regulatory light chain Mrlc promotes contraction (Vicente-Manzanares et al., 2009), while myosin light chain phosphatase (MLCP) protein complex dephosphorylates Mrlc to induce relaxation. MLCP comprises a catalytic subunit Pp1cδ, Mypt1, and a small regulatory subunit, M20 (Ito et al., 2004). Mypt1 enhances MLCP activity and substrate specificity by modulating the catalytic domain conformation (Grassie et al., 2011; Hartshorne et al., 2004). Mypt1 is essential for various developmental processes, including gastrulation, axon guidance, vascular remodelling, and the morphogenesis of the liver, pancreas, and central nervous system (Angulo-Urarte et al., 2018; Barresi et al., 2010; Bremer and Granato, 2016; Dong et al., 2019; Gutzman and Sive, 2010; Huang et al., 2008; Hughes et al., 2020; Kim et al., 2011; Okamoto et al., 2005; Weiser et al., 2009).

In medaka (*Oryzias latipes*), intestinal morphogenesis begins at stage (st.) 21 with medial migration of the endodermal epithelial monolayer to the ventral midline, (Kobayashi et al., 2006). This monolayer then forms a bilayer of mediolaterally elongated cells. In the anterior region, endodermal cells stack and converge at the dorsal midline to form a radial endodermal rod—the intestinal anlage. In contrast, intermediate and posterior gut regions undergo dorsoventral elongation and subsequent cavitation to form a tubular gut. Tube formation proceeds in an anterior-to-posterior sequence, with the anterior and intermediate regions becoming lumenized by st. 25, and the posterior region completing this process by st. 26. During this period, buds of accessory organs such as the liver and swim bladder also emerge (Kobayashi et al., 2006).

In this study, we report a novel medaka mutant identified through N-ethyl-N-nitrosourea (ENU) mutagenesis that develops IA during embryogenesis. Positional cloning identified a nonsense mutation in *mypt1*. Despite normal endodermal migration and gut anlage formation, mutant embryos failed to maintain epithelial continuity in the intermediate intestinal region immediately following lumen formation. This disruption coincided with localized accumulation of phosphorylated Mrlc (pMrlc) and F-actin, consistent with enhanced actomyosin contractility due to loss of Mypt1 function. Our findings establish a genetically defined vertebrate model of Mypt1-associated IA and provide new mechanistic insights into how dysregulated cytoskeletal dynamics compromise epithelial integrity during gut development.

## Results

### The *g1-4* medaka mutant develops IA

The *g1-4* medaka mutant was isolated as a recessive lethal line through our ENU mutagenesis screening for mutations affecting embryonic development and organogenesis.(Yokoi et al., 2007) Examination of homozygous mutant embryos revealed the presence of IA, which developed at st. 32–33, although the penetrance rate of the IA phenotype varied and did not reach 100% (Table 1). In adult medaka, small and large intestines distinguishable by both morphological characteristics and distinct gene expression profiles.(Aghaallaei et al., 2016) However, we could not distinguish small from large intestine in the embryos a either morphologically or by the expression of known molecular markers. Therefore, hereafter, we refer to the region between the liver bud and the cloaca simply as the intestine. Despite of the existence of IA, the remaining intestinal tissue appeared to be properly developed at st. 40, the hatching sage (Fig. 1A, B).

**Fig. 1.**
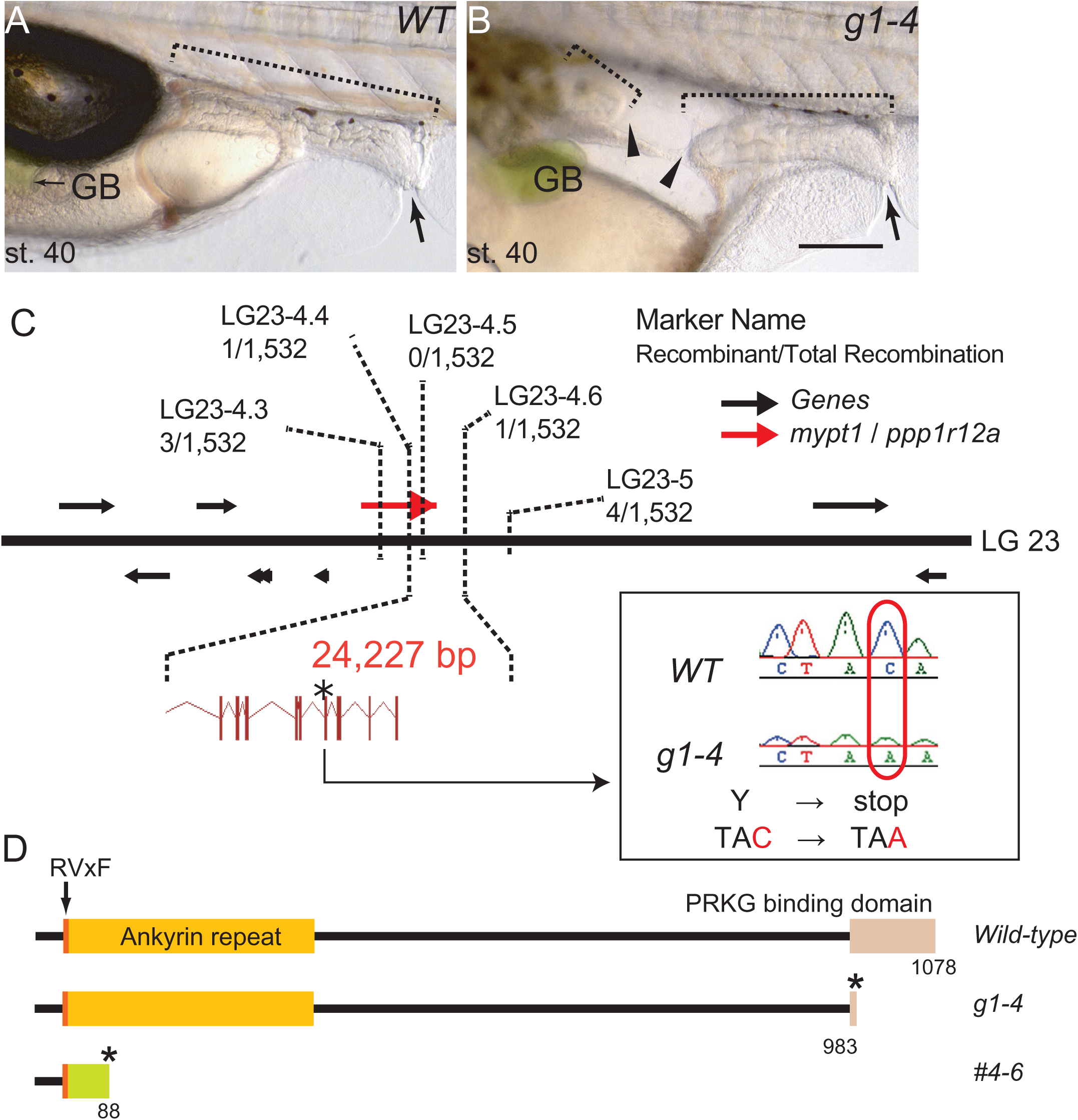
Characterisation of medaka IA mutant, *g1-4*. (A, B) Lateral view**s** of *WT* (A) and *mypt1* mutant (*g1-4*) (B) embryos at st. 40. Note the development of IA in the mutant, represented by the blind ends of the intestine (arrowheads). Anterior is to the left. Arrow, cloaca; GB, gallbladder; Dotted brackets, intestine. Scale bar: 200 µm. **(C)** Schematic presentation of part of LG23. The *g1-4* locus is confined to a 0.13 cM interval between the markers LG23-4.4 and LG23-4.6. Inset: chromatogram of a point mutation in *g1-4*, which gives rise to a premature stop codon. (D) Schematics outlining Mypt1 domains of *WT* (top), *g1-4* (middle), and *mypt1^#4-6^*(bottom) proteins. Asterisks indicate premature translation termination. In *mypt1^#4-6^*, an 11-nucleotide deletion with a 7-nucleotide insertion cause a frame-shift (green), leading to truncation.

**Table 1.** Variation in penetrance among *g1-4* pairs.

| Fish pair |  | IA | TOTAL | % |
| --- | --- | --- | --- | --- |
| Female 1 | Male 1 | 1 | 37 | 2.7 |
| Female 2 | Male 2 | 8 | 45 | 17.8 |
| Female 2 | Male 3 | 8 | 36 | 22.2 |
| Female 3 | Male 2 | 5 | 30 | 16.7 |
| Female 4 | Male 3 | 2 | 20 | 10.0 |
| Female 5 | Male 1 | 4 | 45 | 8.9 |

IA: The number of embryos that develop intestinal atresia.
Total: The number of embryos after deduction of those for which intestinal phenotype could not be determined (dead or abnormal development) .

### The *g1-4* allele encodes *mypt1*

We hypothesized that the inheritance mode of the *g1-4* mutant was recessive, and to identify the affected locus, we performed positional cloning. Initial mapping using M-marker analysis placed the mutation on linkage group 23 (LG23).(Kimura et al., 2004) We then conducted high-resolution linkage analysis using an F2 mapping panel comprising 766 embryos. This analysis narrowed the *g1-4* locus to a 0.13 cM interval between the markers LG23-4.4 and LG23-4.6, corresponding to a genomic region of 24.227 kb (Fig. 1C). Within this region, only one gene, *mypt1*, also known as *ppp1r12a* (*protein phosphatase 1 regulatory subunit 12a*) was identified in the medaka reference genome.

The open reading frame (ORF) of medaka *mypt1* (3,234 bp: LC662536) consists of 25 exons and encodes a predicted protein of 1,078 amino acids that is closely related to human MYPT1 (NP_001137357.1). Medaka Mypt1 has three highly conserved domains, an RVxF motif, an ankyrin repeat, and a protein kinase cGMP-dependent (Prkg) interacting domain (Fig. 1D). Sequencing the *g1-4* allele revealed a C-to-A transversion in exon 22 (C2952A), resulting in a premature stop codon (Fig. 1C, D). The predicted premature termination occurs within the Prkg-interacting domain and eliminates the C-terminal leucine-zipper (LZ) motif, which is essential for interaction with Prkg1α.(Grassie et al., 2011; Surks et al., 1999) As interaction with Prkg1α is essential for actomyosin activation, the C2952A mutation is expected to impair the function of Mypt1.

To confirm whether the *mypt1* mutation causes the IA phenotype observed in the *g1-4* mutants, we tried to rescue the defect using *wild-type* (*WT*) *mypt1* mRNA. However, for unknown reasons, we could not synthesize the full-length *mypt1* mRNA *in vitro*. Alternatively, we generated *mypt1* knockout medaka using the CRISPR/Cas9 system.(Ansai and Kinoshita, 2014) We isolated a mutant line possessing an 11 nucleotide deletion with a 7 nucleotide insertion (*mypt1^#4-6^*, c.127_137delinsTCTGTAT, Fig. 1D, Fig. 2A, B). This mutation creates a transcriptional frame shift that alters 45 codons before introducing a premature stop codon at the 88th codon. Crosses between F1 *mypt1^#4-6^* heterozygotes yielded embryos displaying IA at a frequency of 23.7% (n = 59), and we successfully established a stable mutant line. In the F2 generation, the penetrance of the IA phenotype varied among sibling, similar to the original *g1-4* mutant line (Table 2), with approximately 15–30% of total siblings exhibiting IA, - higher penetrance than in the original mutant. For further phenotypic analysis, we used this newly generated mutant line, *mypt1^#4-6^*. We could also see dilation of the intestine upstream of the atretic region at the hutching stage, a hallmark commonly associated with IA in *mypt1^#4-6^* embryos (Fig. 3A–D). Taken together, we concluded that *mypt1* mutation is responsible for the IA phenotype.

**Fig. 2.**
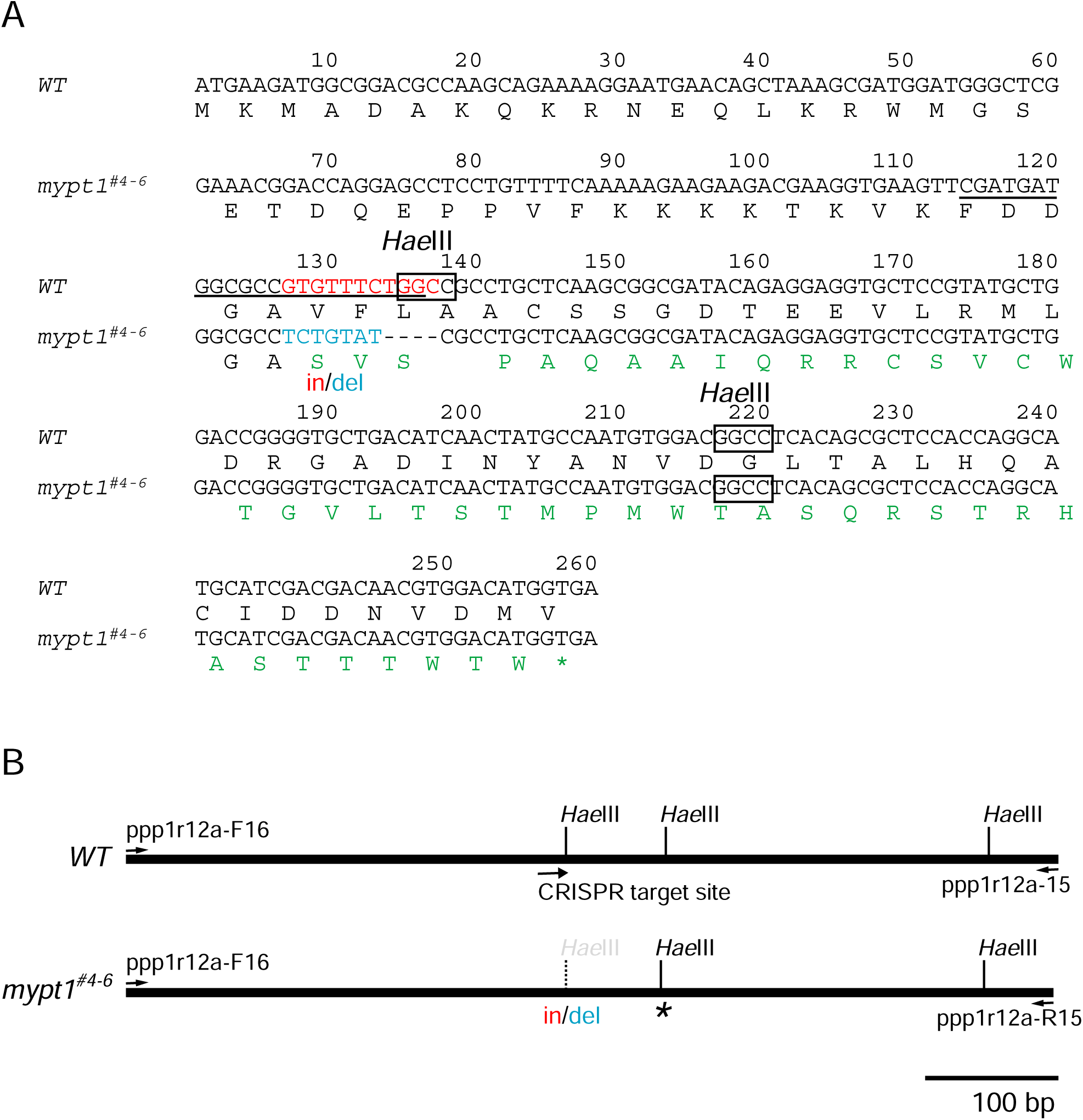
Comparison of *WT* and *mypt1^#4-6^* allele sequences. (A) The 11-nucleotide deletion and 7 nucleotide insertion are shown in red and blue, respectively. The CRISPR target site is underlined. The predicted amino acid sequence resulting from the frame shift is shown in green. Asterisk indicates premature stop codon. *Hae*III sites (GGCC) that are used for genotyping by restriction fragment length polymorphism (RFLP) assays are indicated. (B) Schematic outline of PCR products amplified from *WT* and *mypt1^#4-6^* alleles. Primer positions (*ppp1r12a*-F15 and *ppp1r12a*-R16), the CRISPR target site and *Hae*III sites are indicated. The *Hae*III site lost in *mypt1^#4-6^* is shown in grey. The introduced stop codon in *mypt1^#4-6^* is marked by an asterisk.

**Fig. 3.**
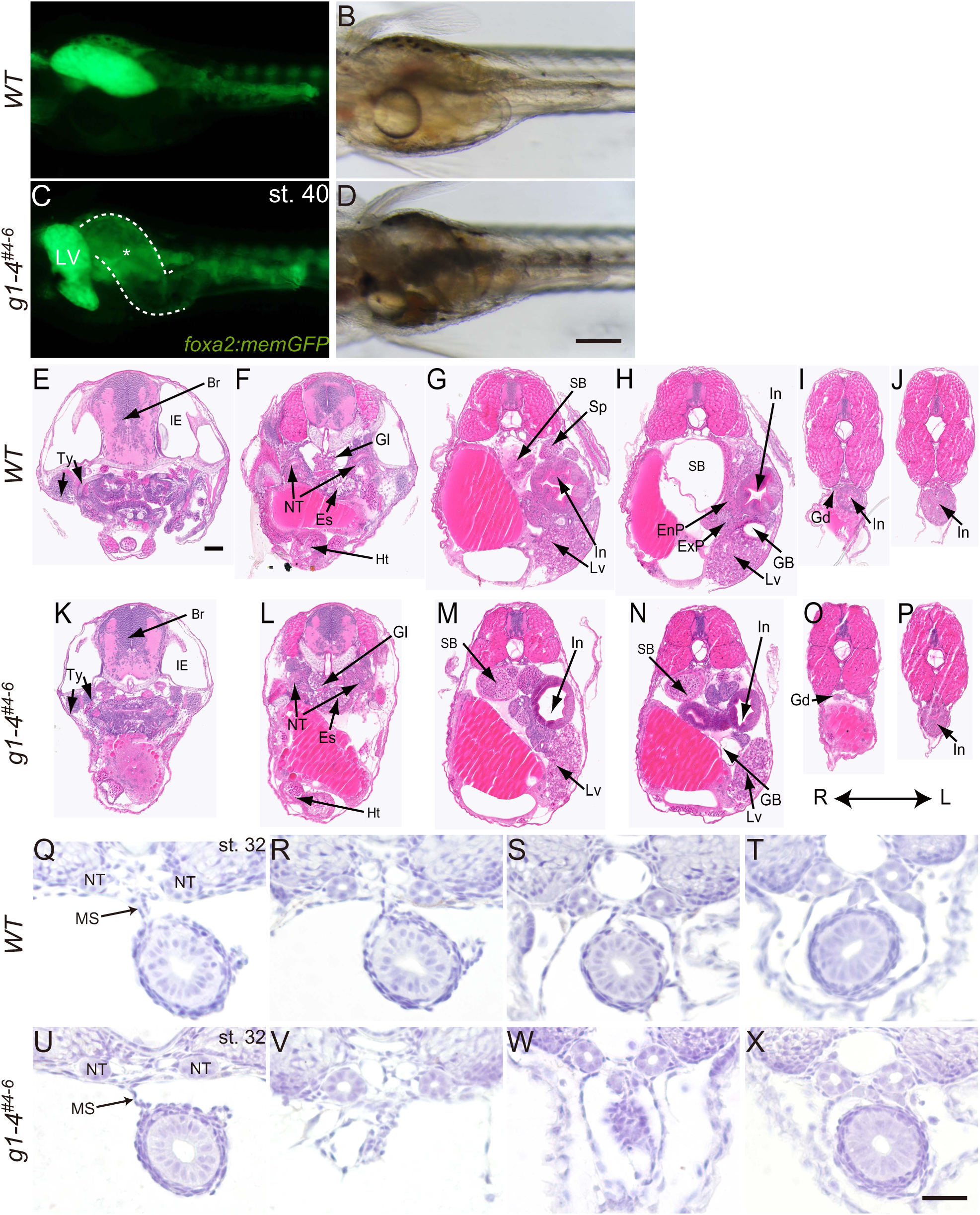
Organogenesis proceeds normally in *mypt1* mutants except for the presence of IA. (A–D) Dilation of the intestine in *mypt1* mutants. Ventral views of *WT* (A, C) and *mypt1#4-6* (B, D) embryos at st. 40 in the Tg[*foxa2:memGFP*] medaka line. GFP fluorescence and brightfield images are shown in (A, B) and (C, D), respectively. White dotted lines outline the intestine. Note the significant dilation anterior to the IA lesion in the mutant (asterisk). LV, liver. Scale bar: 200 µm. (E–P) Histological sections of st. 40 embryos. *WT*: (E–J); *mypt1* mutant: (K–P). Panels show comparable anatomical levels: (E and K), (F and L), (G and M), (H and N), (I and O), and (J and P). Abbreviations: EnP, endocrine pancreas; Es, esophagus; ExP, exocrine pancreas; GB, gall bladder; Gd, gonad; Gl, glomerulus; Ht, heart; IE, inner ear; In, intestine; Lv, liver; NT, nephric tubule; Sp, spleen; AB, swim bladder; Ty, thyroid; R, right; L, left. Scale bar: 50 µm. (Q–X) Histological micrographs of the intestine in *WT* (Q–T) and *mypt1^#4-6^* (U–X) embryos at st. 32, stained with haematoxylin. (U) Region anterior to IA; (V) IA lesion; (W) anterior blind-end of posterior intestine; (X) region posterior to the IA. *WT* sections (Q–T) correspond anatomically to (U–X). Note normal morphology in (U) and (X). MS, mesentery; NT, nephric tubule. Scale bar: 20 µm.

**Table 2.**
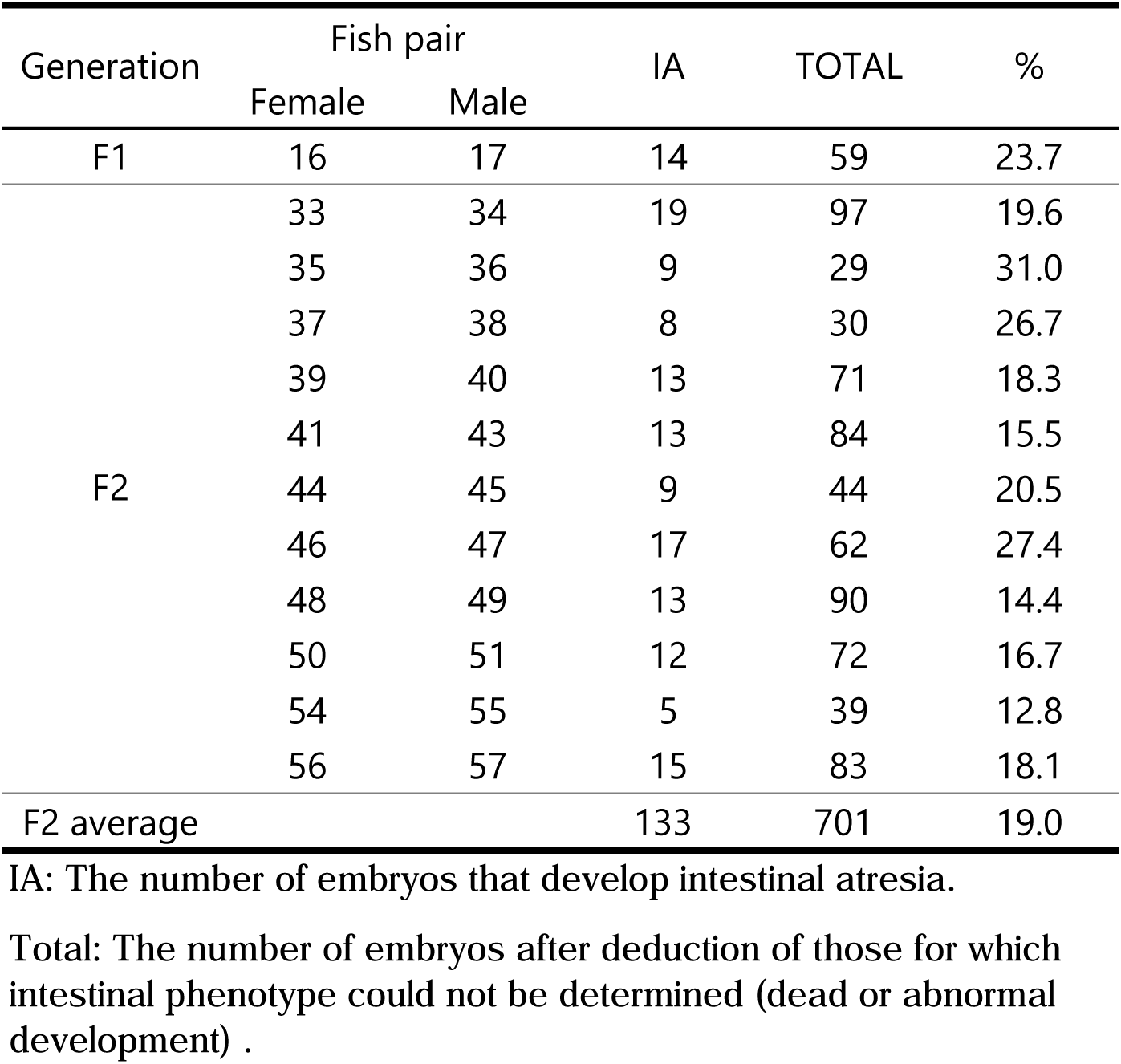
Variation in penetrance among *mypt1 ^#4-6^* pairs.

### Intestinal Development Is Largely Preserved in *mypt1#4-6* Mutants, Except at the Atresia Lesion

To investigate the phenotype of the *mypt1^#4-6^* mutant, we performed histological analyses on st. 40 larvae, immediately after hatching. Serial cross-sections were prepared from *WT* (*n* = 3) and *mypt1^#4-6^*mutant larvae (*n* = 3) and compared. The mutants exhibited intestinal atresia, whereas other organs appeared to develop normally (Fig. 3E–P). To further assess the epithelial architecture at earlier stages, we performed conventional histological analysis using plastic sections at st. 32. These sections showed that the embryonic intestine appeared morphologically intact in *mypt1^#4-6^* mutants, except at the atretic lesion. The mesenchymal cells, which give rise to SM and connective tissue, surrounded the endodermal epithelium similarly to *WT*, and this organization was also preserved in unaffected regions of the mutant intestine (Fig. 3Q–X) (Wallace et al., 2005a). Homozygous mutants were generally lethal, with only few individuals surviving to adulthood. Most homozygous mutants die after hatching. Thus, intestinal atresia is very likely one of the major causes for lethality, as affected larvae are unable to feed after hatching. However, we cannot rule out the possibility that other factors contribute to lethality. Importantly, while most survivors exhibited poor health, some were indistinguishable from *WT* fish (data not shown).

Cytokeratin, a marker of epithelial cells and intermediate filament protein, predominantly localises to the apical plasma membrane in polarized epithelia (Fig. 4A) (Yang et al., 2020). Consistent with this, the intestinal epithelium of st. 31 *WT* embryos showed strong apical localization of cytokeratin (Fig. 4Bb). In *mypt1^#4-6^* mutants, the intestinal epithelium— excluding the atretic lesion—displayed cytokeratin distribution comparable to *WT* (Fig. 4Cb).

**Fig. 4.**
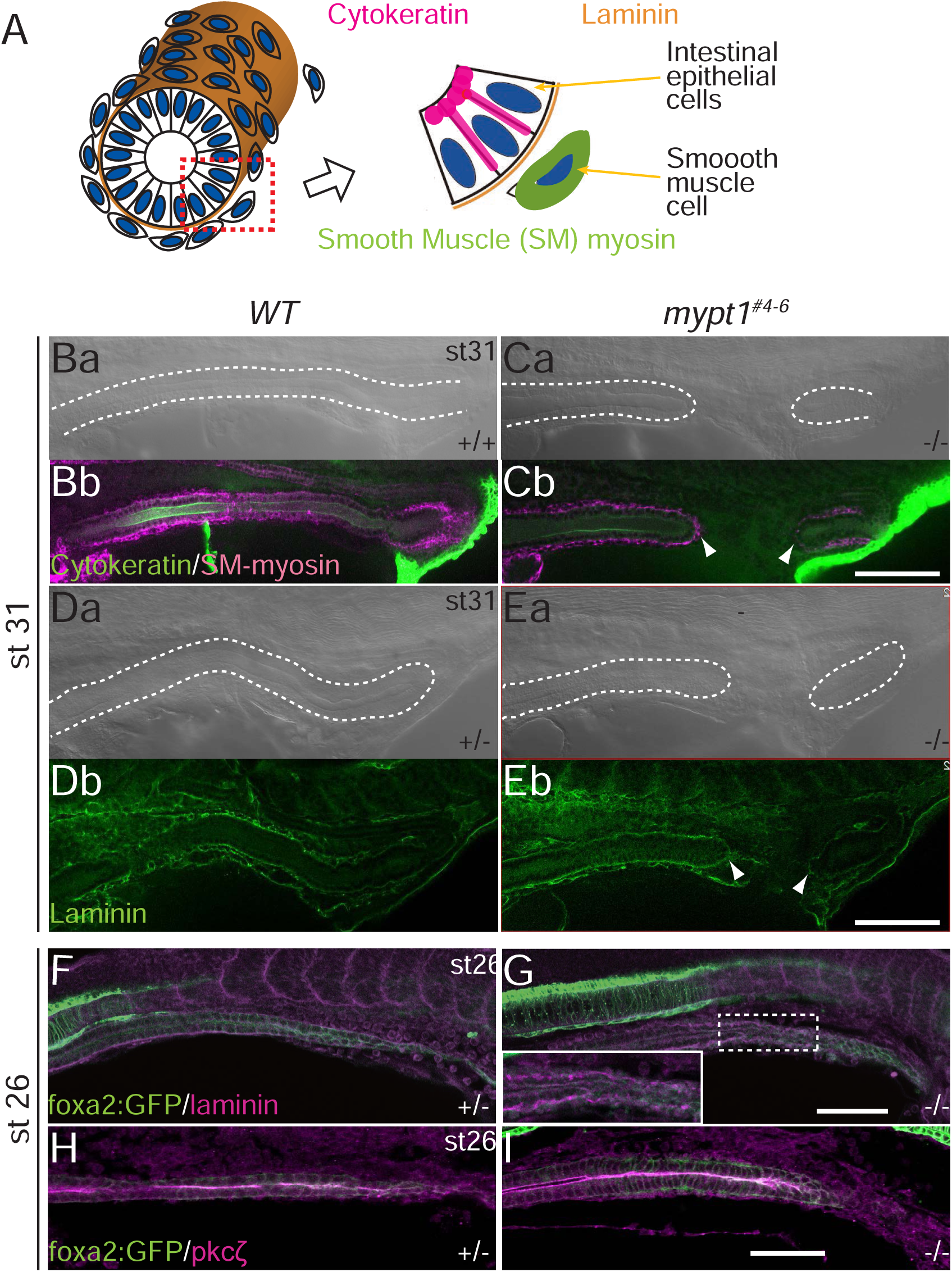
Apico-Basolateral polarity is maintained in *mypt1* mutants. (A) Schematic diagram of the distribution of cytokeratin, laminin and SM myosin in the developing intestine. (B–I) Whole mount immunofluorescence micrographs of cytokeratin and SM myosin (B, C), laminin (D–G) and Pkcζ (H, I) in *WT* (B, D, F, H) and *mypt1^#4-^*(C, E, G and I) as left lateral view. White dotted lines mark the outline of intestine in a differential interference contrast (DIC) micrograph corresponding to a fluorescence image. Note that basement membrane was fragmented in the mutant embryos at st. 26 before IA onset (F, n = 11; G, n=9). (H, I) Immunohistochemistry for Pkcζ in long-axis optical sections of st. 26 embryo intestines. The epithelium of the intestine was visualized with *Tg*[*foxa2:memGFP*] (H, n = 9; I, n = 10). Note that the apical localization of Pkcζ was not disturbed in *mypt1* mutants (I). Arrowheads, blind-ends of IA. Scale bars: 50 µm.

SM cells, labelled by SM-specific myosin (SM-myosin), properly surrounded the intestinal epithelium in both *WT* and *mypt1^#4-6^* embryos at st. 31 (Fig. 4Ba–Cb). Notably, the blind end of the atresia lesion in mutants was also lined by SM. Furthermore, the basement membrane, labelled by laminin, was present beneath the epithelium even at the blind-end of the atresia in the mutant (Fig. 4Da–Eb). These findings indicate that the fundamental structure of the gut—including epithelial apico-basolateral polarity, SM layer, and basement membrane—is preserved normally in the region outside the atresia. This is consistent with observations in human IA cases.

To elucidate the sequence of events that lead to atresia formation in *mypt1^#4-6^* embryos, we carefully monitored live embryos but could not detect morphological signs of IA until st. 28. We first examined endodermal development, as the endoderm gives rise to the intestine, to determine whether endodermal defects were present before or during intestinal tube formation. Expression of *foxa2*, a marker of the endoderm and its derivatives including the intestine (Kobayashi et al., 2006), was indistinguishable between *WT* and *mypt1^#4-6^* embryos up to st. 25 (Fig. 5), suggesting that early endoderm formation and migration are not disrupted in the mutant.

**Fig. 5.**
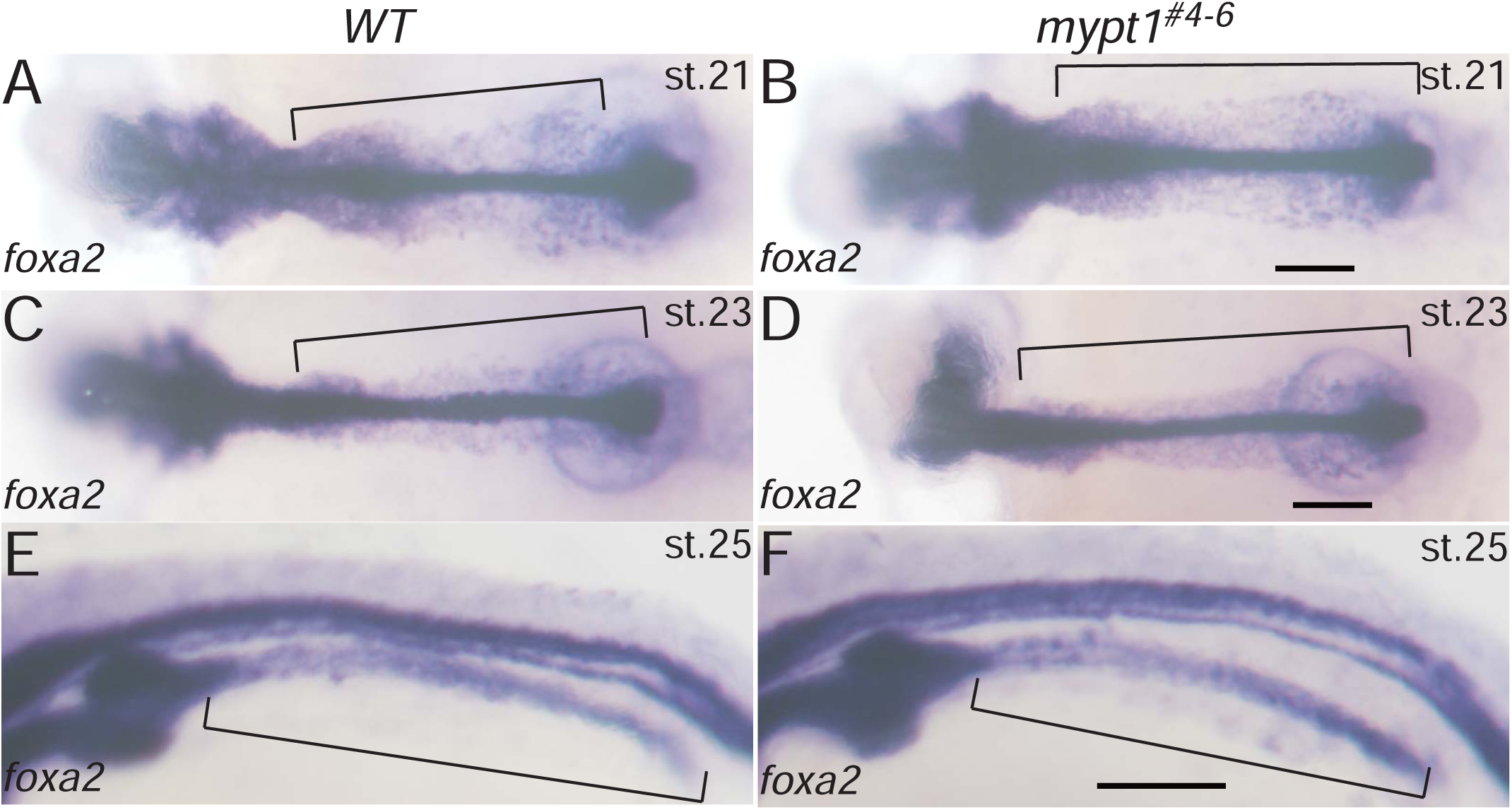
Endodermal migration to form the intestinal anlage is intact in *mypt1* mutants. *WT* (A, C, E, n = 10) and *mypt1* mutant (B, D, F, n = 10) embryos were subjected to whole mount *in situ* hybridization for *foxa2*. Dorsal (A–D) and dorsolateral views (E–F) are shown. Embryonic stages are indicated in the top right. Brackets mark endoderm and intestinal anlage. Scale bar: 100 µm.

Importantly, at st. 26—when intestinal tube formation is completed (Kobayashi et al., 2006)—fragmentation of the basement membrane was observed in 8 out of 9 *mypt1*#4-6 mutants (Fig. 4G), whereas none of the 11 *WT* embryos showed such fragmentation (Fig. 4F). These results indicate that molecular events leading to IA had already begun by st. 26, even though no obvious morphological anomalies were yet visible.

To investigate whether epithelial disintegration due to apoptosis or altered proliferation contributed to basement membrane fragmentation, we performed acridine orange staining at st. 27. This revealed no detectable apoptotic cells in either *WT* or *mypt1^#4-6^* intestines (Fig. 6A, B). TUNEL assays also showed no significant difference between *WT* and mutant embryos (data not shown). Furthermore, proliferation, assessed by anti-phospho-histone H3 staining, showed no significant difference in the number of mitotic cells between *WT* and mutant intestines (Fig. 6E–I). These data suggest that neither apoptosis nor altered proliferation underlies the basement membrane fragmentation and IA in the mutant.

**Fig. 6.**
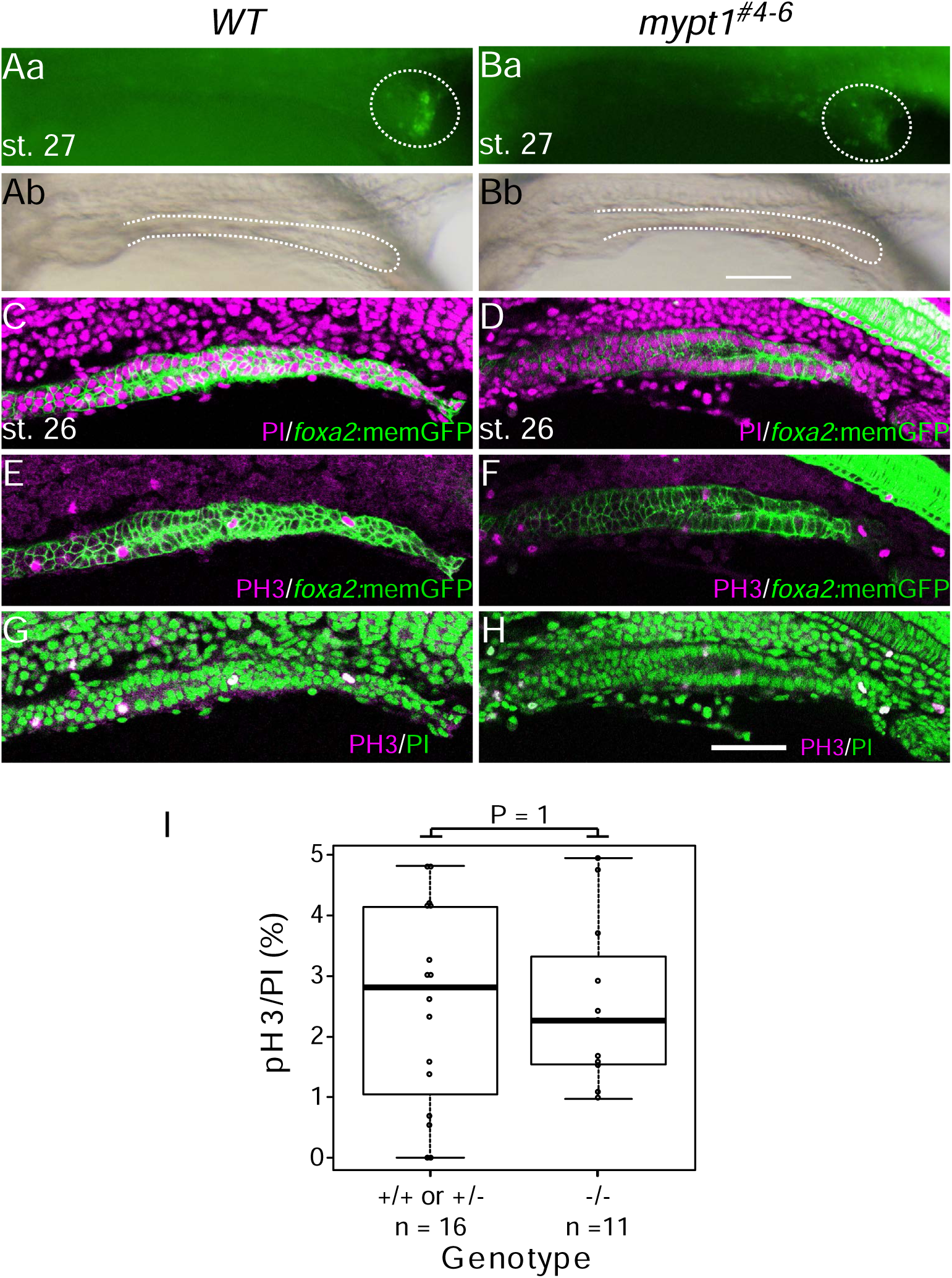
Cell death and cell proliferation are not involved in IA development. (A–H) Cell death and cell proliferation in the intestine are not significantly altered in *mypt1* mutants. (A, B) Acridine orange staining of *WT* embryos (Aa, *n* = 12) and *mypt1* mutants (Ba, *n* = 5) at st. 27. (Ab, Bb) Brightfield images corresponding to (Aa) and (Ba), respectively. Dotted ellipses mark apoptotic cells surrounding the cloacal opening, where apoptosis is normally observed (Parkin et al., 2009). (C–H) *WT* (C, E, G) and *mypt1* mutant (D, F, H) embryos at st. 26 were stained with anti-phosphorylated histone H3 (PH3) antibody and/or propidium iodide (PI). (I) Quantification of PH3-positive cell ratio to total cell number in the intestinal region was not significantly different between *WT* (*n* = 16) and *mypt1* mutants (*n* = 11; chi-square test, *P* = 1). Scale bar: 50 µm.

Although apico-basolateral polarity was preserved in the gut epithelium, localised epithelial-mesenchymal transition (EMT) could hypothetically result in loss of polarity and basement membrane fragmentation, as occurs during gastrulation (Savagner, 2015). To explore this possibility, we examined the expression of EMT-inducing transcription factors, *snai1a*, *snai1b*, and *snai2* (Scheibner et al., 2021), via whole-mount in situ hybridization (WISH). None of these genes were expressed in the intestine of either *WT* or mutant embryos at st. 26 (Fig. 7A–F). Consistently, the subcellular localization of Pkcζ, an apical epithelial marker, was not disrupted in *mypt1*#4-6 intestines (Fig. 4H–I). Taken together, these observations indicate that basement membrane fragmentation and IA formation in *mypt1^#4-6^* embryos do not result from EMT or loss of epithelial polarity.

**Fig. 7.**
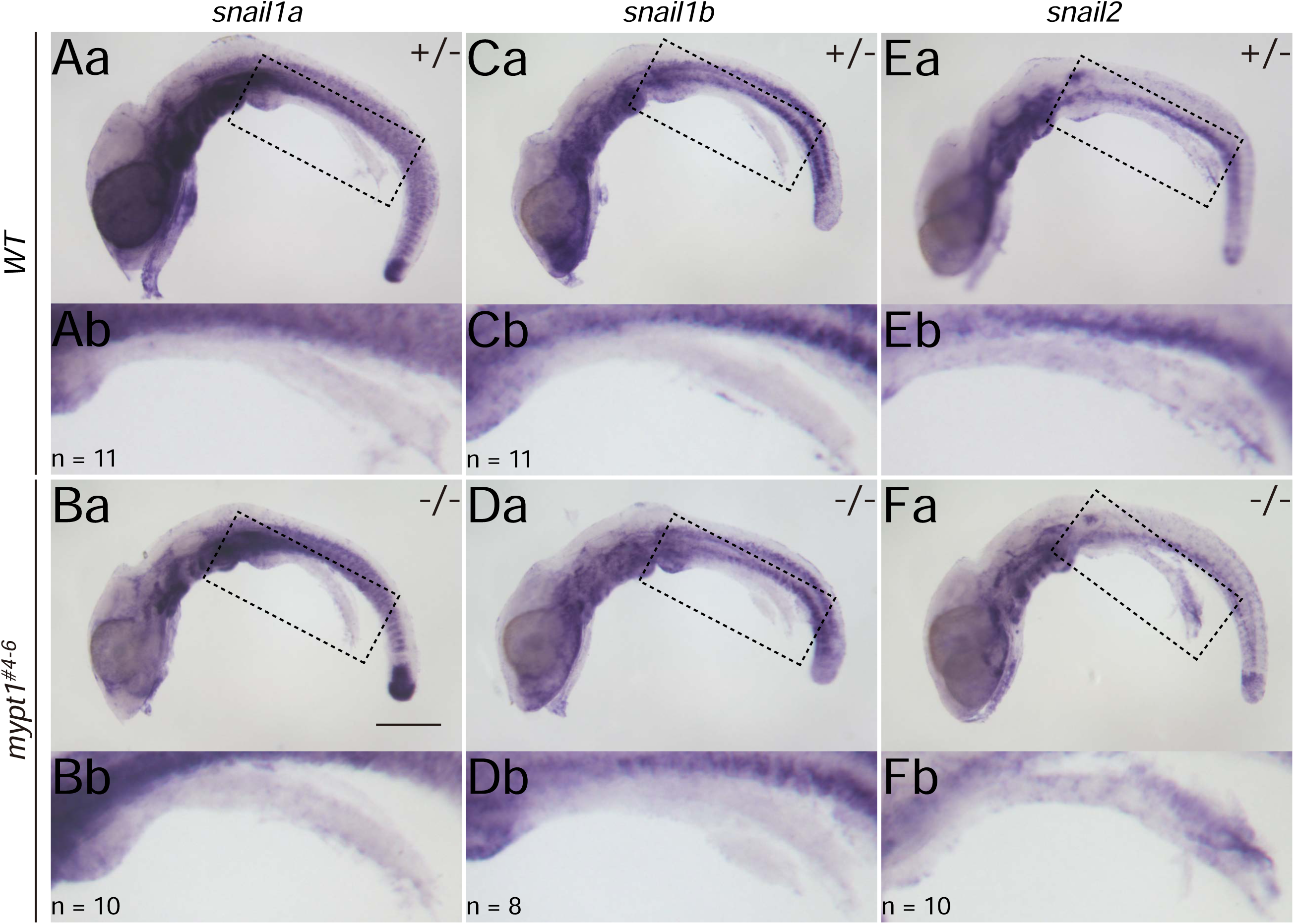
Epithelial–mesenchymal transition (EMT) is not involved in IA formation in *mypt1* mutants. Lateral views of st. 26 embryos subjected to whole mount *in situ* hybridization to detect EMT markers: WT (A, C, E); mutants (B, D, F). (A, B) for *snai1a*, (C, D) for *snai1b*, and (E, F) for *snai2*. To enhance visibility, the intestine was detached from the dorsal mesentery. Dotted boxes in (Aa-Fa) are enlarged in (Ab-Fb), respectively. The number of samples examined is indicated in the bottom left corner. Scale bar: 200 µm.

### Mypt Family Members Are Unlikely to Compensate for the Loss of MYPT1 in a Tissue-Specific Manner

In zebrafish, *mypt1* mutations cause a spectrum of morphogenetic abnormalities, including defects in liver and pancreas development, dilation of brain ventricles, and mispositioning of motoneurons (Bremer and Granato, 2016; Dong et al., 2019; Gutzman and Sive, 2010; Huang et al., 2008). In contrast, these phenotypes were not observed in *mypt1* mutant medaka, suggesting possible species-specific differences in gene function or redundancy within *mypt* gene family. To explore whether this phenotypic divergence could be explained by differential expression or functional compensation by other *mypt* family members, we identified *mypt* homologs in the medaka genome and analysed their spatial expression patterns by WISH.

In humans, the myosin phosphatase regulatory subunit family consists of *MYPT1* (PPP1R12A), *MYPT2* (PPP1R12B), and *MBS85* (PPP1R12C) (Gutzman and Sive, 2010; Huang et al., 2008). To determine whether other *mypt* family genes might compensate for the loss of *Mypt1* function in tissues other than the intestine in medaka mutants, we searched the medaka genome and identified three members of the *mypt* family: *mypt1*, *mbs85*, and a novel gene that diverges from the ancestral lineage shared by *mypt1* and *mypt2*, which we designated *mypt1/2-related* (*mypt1/2-r*) (Table 3; Fig. 8). Notably, no clear *mypt2* ortholog was found in the medaka genome. Although medaka possesses a gene annotated as *mypt3*, this gene belongs to the *Ppp1r16* family rather than the *Ppp1r12* family and therefore was excluded from further analysis.

**Fig. 8.**
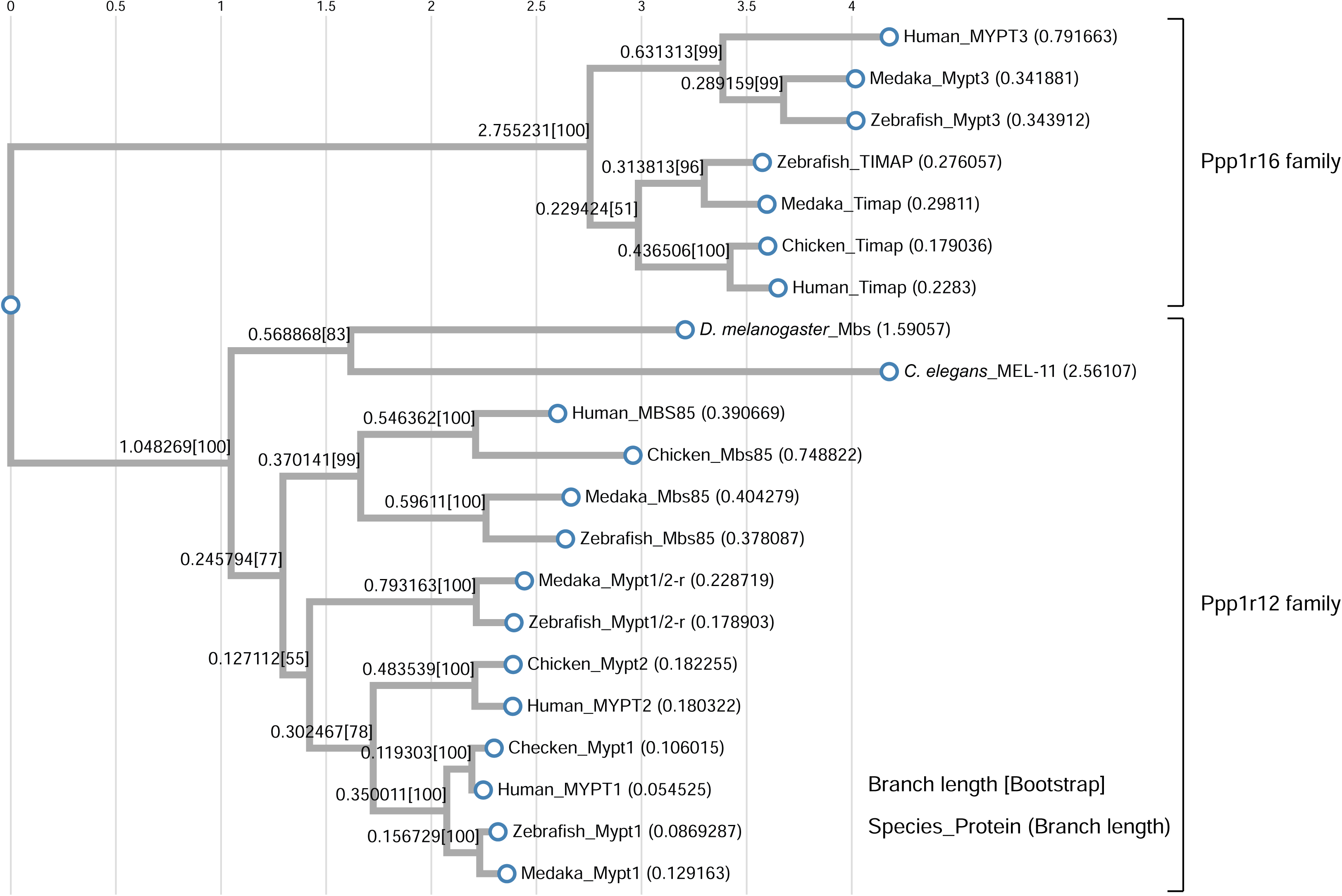
Phylogenetic analysis of the *Mypt* gene family. Phylogenetic tree conducted from protein sequences of Mypt family genes listed in Table 3.

**Table 3.** List of protein sequences used for ClustalW analysis.

| **Species** | **Protein Name** | **Accession Number** |
| --- | --- | --- |
| *Homo sapiens* | MYPT1 | NP_002471.1 |
| *Gallus gallus* | Mypt1 | NP_990454.1 |
| *Danio rerio* | Mypt1 | NP_001003870.2 |
| *Oryzias latipes* | Mypt1 | XP_023807904.1 |
| *Homo sapiens* | MYPT2 | NP_002472.2 |
| *Gallus gallus* | Mypt2 | XP_040508700.1 |
| *Homo sapiens* | MBS85 | NP_060077.1 |
| *Gallus gallus* | Mbs85 | XP_025002065.1 |
| *Danio rerio* | Mbs85 | NP_001071047.1 |
| *Oryzias latipes* | Mbs85 | XP_023810293.1 |
| *Danio rerio* | Mypt1/2-r | XP_002666820.2 |
| *Oryzias latipes* | Mypt1/2-r | XP_020559658.2 |
| *Homo sapiens* | MYPT3 | NP_001316373.1 |
| *Danio rerio* | Mypt3 | XP_001334473.1 |
| *Oryzias latipes* | Mypt3 | XP_011487151.1 |
| *Homo sapiens* | TIMAP | NP_056383.1 |
| *Gallus gallus* | Timap | NP_001026022.2 |
| *Danio rerio* | Timap | XP_001340092.6 |
| *Oryzias latipes* | Timap | XP_004068819.1 |
| *D. melanogaster* | mbs | NP_001261919.1 |
| *C. elegans* | MEL-11 | AAB47273.1 |

We examined the expression patterns of *mypt1*, *mypt1/2-r*, and *mbs85* by WISH. Both *mypt1* and *mypt1/2-r* were ubiquitously expressed at low levels, with slightly elevated expression in the head region. *mbs85* exhibited lower overall expression than the other two genes but displayed a similar spatial pattern. No significant differential expression was detected in the developing intestine (Fig. 9A–F). RT-PCR analysis of dissected head and intestinal tissues confirmed the expression of all three genes, supporting their broad, low-level distribution. These findings suggest that the intestinal specificity of the atresia phenotype in *mypt1* mutants is not due to tissue-specific expression of *mypt* family genes. Furthermore, given the ubiquitous—albeit low—expression of these genes, it is possible that the phenotypes observed in the zebrafish *mypt1* mutant were not reproduced in the medaka mutant due to functional compensation by related genes.

**Fig. 9.**
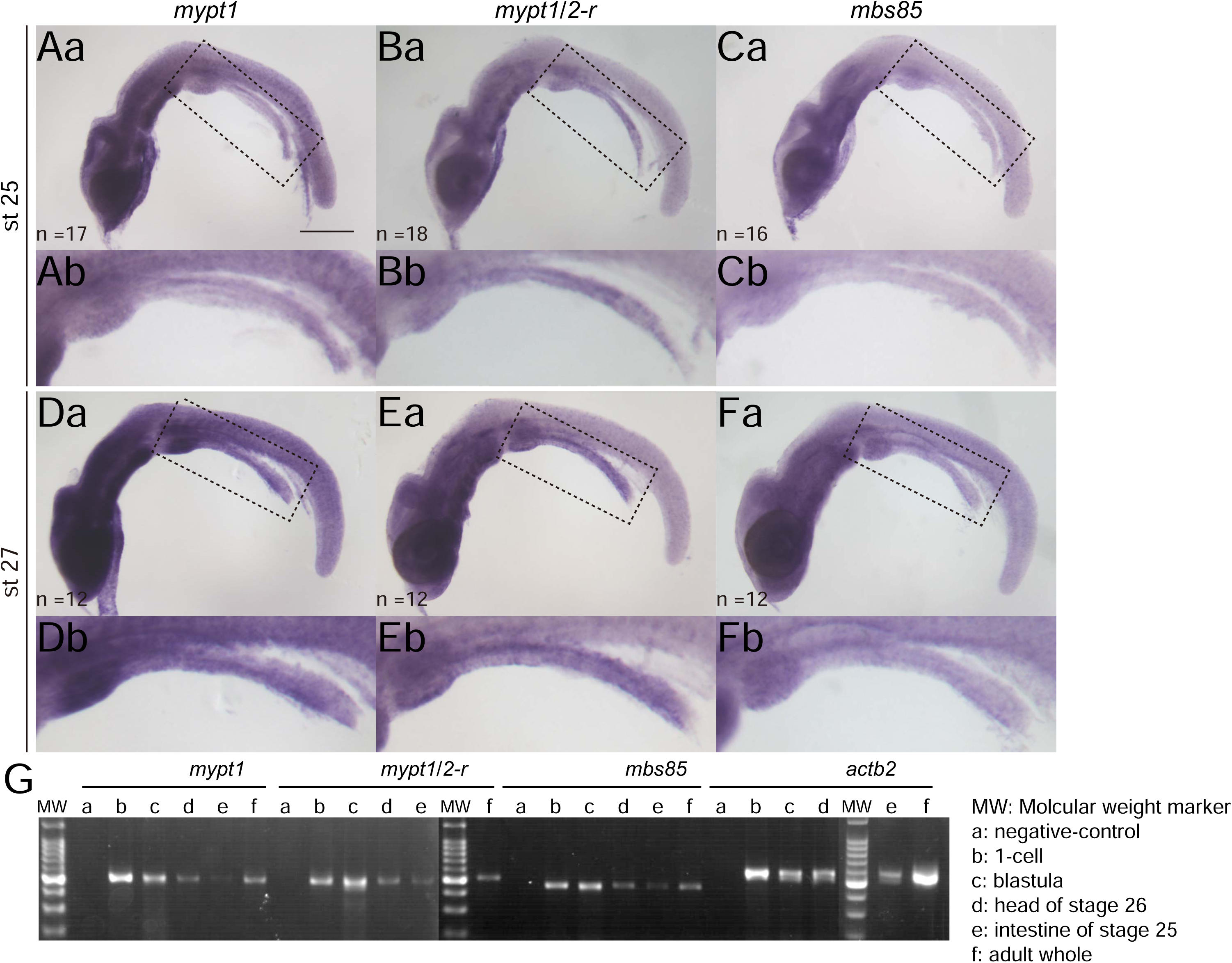
Expression patterns of *Mypt1* family genes. (A–F) Whole-mount *in situ* hybridization of *mypt1* (A, D), *mypt1/2-r* (B, E), and *mbs85* (C, F). (A–C) st. 25; (D–F) of st. 27 in *WT*. Dotted boxes in (Aa-Fa) are enlarged in (Ab-Fb), respectively. The number of samples examined is indicated in the bottom left corner. Scale bar: 200 µm. MW: Molecular weight marker (DM2100, SMBIO, Taiwan) (G) RT-PCR analysis. PCRs using first-strand cDNA from: (a) negative control, (b) 1-cell stage, (c) blastula stage, (d) head region at stage 26, (e) intestine region at stage 25, and (f) adult whole body.

### Mypt1 Is Maternally Expressed

Previous studies in zebrafish have shown that maternal knockdown of *mypt1* using translation-blocking morpholinos leads to convergent extension (CE) defects during gastrulation (Weiser et al., 2009). In contrast, such defects were not observed in the *mypt1^#4-6^* mutant medaka line. To investigate whether maternal *mypt1* transcripts contribute to early embryogenesis and mask early developmental phenotypes, we performed RT-PCR on one-cell stage embryos. The results confirmed the presence of *mypt1* transcripts at this stage, indicating that maternal *mypt1* mRNA is supplied during oogenesis. This maternal expression is likely to compensate for the absence of zygotic *mypt1*, thereby preventing CE defects in *mypt1^#4-6^*mutants (Fig. 9G).

### Actomyosin Regulation Is Perturbed in *mypt1* Mutant Intestine

Phosphorylation of myosin regulatory light chain (Mrlc) governs the contractility of non-muscle myosin II (NMII). (Ito et al., 2004) Dephosphorylated Mrlc reduces NMII activity and cytoskeletal contractility, while pMrlc activates NMII. Thus, NMII-driven contractility is determined by the balance between phosphorylation by myosin light chain kinase (Mlck) and dephosphorylation by myosin light chain phosphatase (MLCP). *Mypt1* encodes the regulatory subunit of MLCP that is essential for its enzymatic function. Therefore, in *mypt1* mutants, elevated levels of pMrlc are expected.

In *WT* embryos, pMrlc exhibited weak and diffuse localisation specifically on both the apical and basal surfaces of intestinal epithelial cells (Fig. 10A, st. 25; 10C, st. 27). As predicted, *mypt1* mutants displayed increased pMrlc levels at st. 25 compared with *WT* embryos (Fig. 10B). This difference was not apparent at st. 24 (data not shown). Notably, pMrlc was not uniformly increased along the intestinal axis but appeared enriched in regions corresponding to presumptive intestinal atresia (IA) lesions (Fig. 10B). Higher magnification revealed that this increased pMrlc signal localized specifically to the intestinal epithelium, as marked by *foxa2:memGFP* expression (Fig. 10Bd). By st. 27, this abnormal expression pattern became more pronounced (Fig. 10D), and in some embryos, pMrlc was strongly expressed along the entire intestinal tract (Fig. 10E; 2 out of 12 samples). These findings indicate that pMrlc regulation is markedly altered in *mypt1* mutants from st. 25 onward, particularly in regions prone to IA formation (Fig. 10F).

**Fig. 10.**
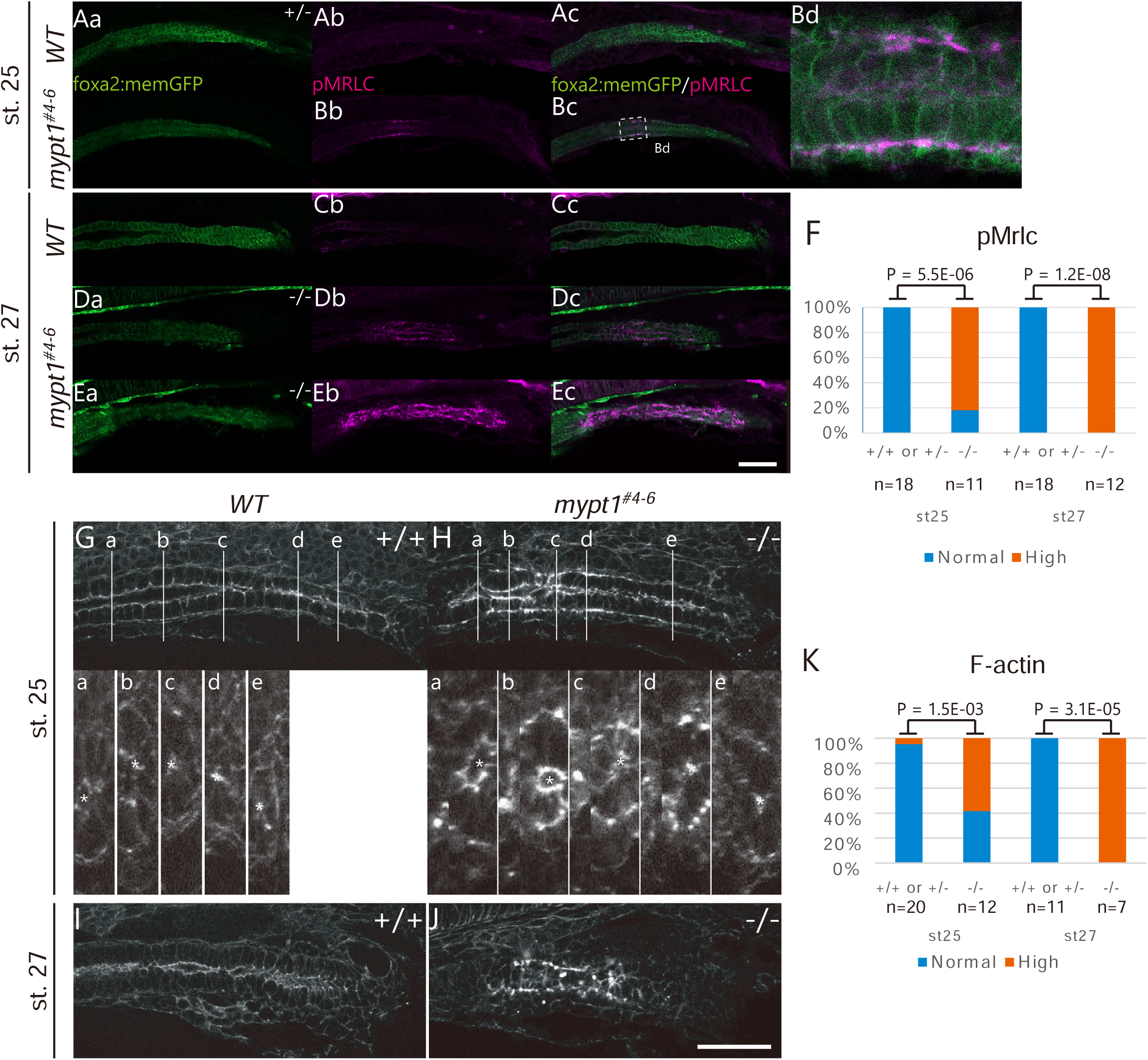
Actomyosin is abnormally activated in the developing intestine of *mypt1* mutants. (A**–**E) Fluorescence micrographs of pMrlc for *WT* (A, C) and *mypt1^#4-6^*(B, D, E). st. 25 embryos are shown in A and B, whereas st. 27 embryos are shown in C, D and E. Note significant accumulation of pMrlc in mutant intestinal epithelium (Bb, Db, Eb). Dotted boxes in Bc is enlarged in Bd. Scale bar, 50 µm. (F) The proportion of embryos possessing an abnormal increase of pMrlc in embryonic intestine (Fisher’s exact test). (G-J) Fluorescence micrographs of F-actin stained with phalloidin. (a**–**e) represent short axis sections of intestine. *WT* (G, I) and *mypt1****^#4-6^*** (H, J) st.25 (G, H) and st. 27 (I, J) embryos are shown. (H, J) Note abnormal accumulation of F-actin in the *mypt1* mutant intestine. Scale bar: 50 µm. *, an intestinal lumen. (K) The proportion of embryos possessing an abnormal increase of F-actin in embryonic intestine (Fisher’s exact test).

Because pMrlc binds NMII and promotes actin-myosin interaction, increased and mis-localised pMrlc is likely to enhance actomyosin contractility and redistribute mechanical forces (Vicente-Manzanares et al., 2009). In *WT* intestinal epithelial cells, F-actin was localised to the cortical region beneath the plasma membrane and was slightly enriched at the apical surface (Fig. 10G and I). In contrast, although intestinal lumen formation appeared morphologically similar between *WT* and *mypt1* mutants, F-actin accumulation was elevated in both the apical and basal cortical regions of *mypt1* mutant cells (Fig. 10H, J, and K). This phenotype is consistent with previous reports showing that increased pMrlc enhances actin filament assembly (Gutzman and Sive, 2010), whereas reduced pMrlc leads to decreased F-actin levels.(Bavaria et al., 2011; Ray et al., 2007) Together, these results indicate that loss of *mypt1* function leads to upregulation of pMrlc, suggesting hyperactive actomyosin contractility of intestinal epithelium. This aberrant mechanical activity may compromise epithelial integrity and contribute to IA development in *mypt1* mutant embryos.

### Smooth Muscle Myosin Function Partially Contributes to the Development of Intestinal Atresia (IA)

Given that Mypt1 regulates SM myosin activity, abnormalities in the SM layer surrounding the intestinal epithelium may play a role in the pathogenesis of intestinal atresia (IA). In the medaka genome, we identified two homologs of *MYH11*: *myh11a* on linkage group (LG) 1 and *myh11b* on LG 8. However, RT-PCR analysis at 2 days post-fertilization (dpf) revealed expression of *myh11a* only (data not shown), prompting us to focus our investigation on *myh11a*.

To disrupt *myh11a* function, we designed three overlapping single-guide RNAs (sgRNAs) targeting the ATP-binding domain of *myh11a* (Fig. 11A). These sgRNAs were injected into medaka embryos at the 1–2 cell stage, either alone or in combination with Cas9 nuclease. To confirm the effect of CRISPR-mediated knock-out, we performed immunohistochemical analysis of SM myosin expression at a developmental stage when the SM layer is well established. In the embryos injected without Cas9 at 6 dpf, SM myosin was robustly detected in the smooth muscle layer surrounding the intestinal epithelium (Fig. 11B; 6/6 embryos). In contrast, embryos co-injected with Cas9 exhibited either a complete loss of SM myosin expression (Fig. 11C; 2/6 embryos) or an aberrant, patchy expression pattern (Fig. 11D; 4/6 embryos), indicating effective disruption of functional SM myosin. We then assessed whether IA still occurred under these SM myosin-deficient conditions in *mypt1^#4-6^* homozygous mutant embryos. Notably, IA was absent in 4 out of 10 *mypt1^#4-6^* embryos co-injected with *myh11a* sgRNAs and Cas9, whereas all 9 embryos injected with *myh11a* sgRNAs alone (without Cas9) developed IA (Table 4). To further validate the role of *myh11a*, we performed a morpholino (Mo)-mediated knockdown using a splice-blocking morpholino targeting the exon 2–intron 2 junction (Mo-*myh11a*-E2I2; Fig. 11E). RT-PCR confirmed efficient disruption of normal splicing (Fig. 11F), and partial rescue of IA was observed in morpholino-treated embryos (Table 5). These findings suggest that *myh11a*-dependent SM function contributes to IA pathogenesis. However, as immunohistochemical signal of Myh11a is not detected around the region where IA occur at st. 28 when IA becomes morphologically detectable, the SM contraction is not likely involved in IA formation (data not shown). Non-smooth muscle expression of *myh11a* (e.g., in intestinal epithelial cells), or alternatively, cells expressing *myh11a* located distant from the intestinal epithelium, may contribute non-cell-autonomously to epithelial integrity at early stages in medaka, but likely play only a partial, not primary, role (see Discussion).

**Fig. 11.**
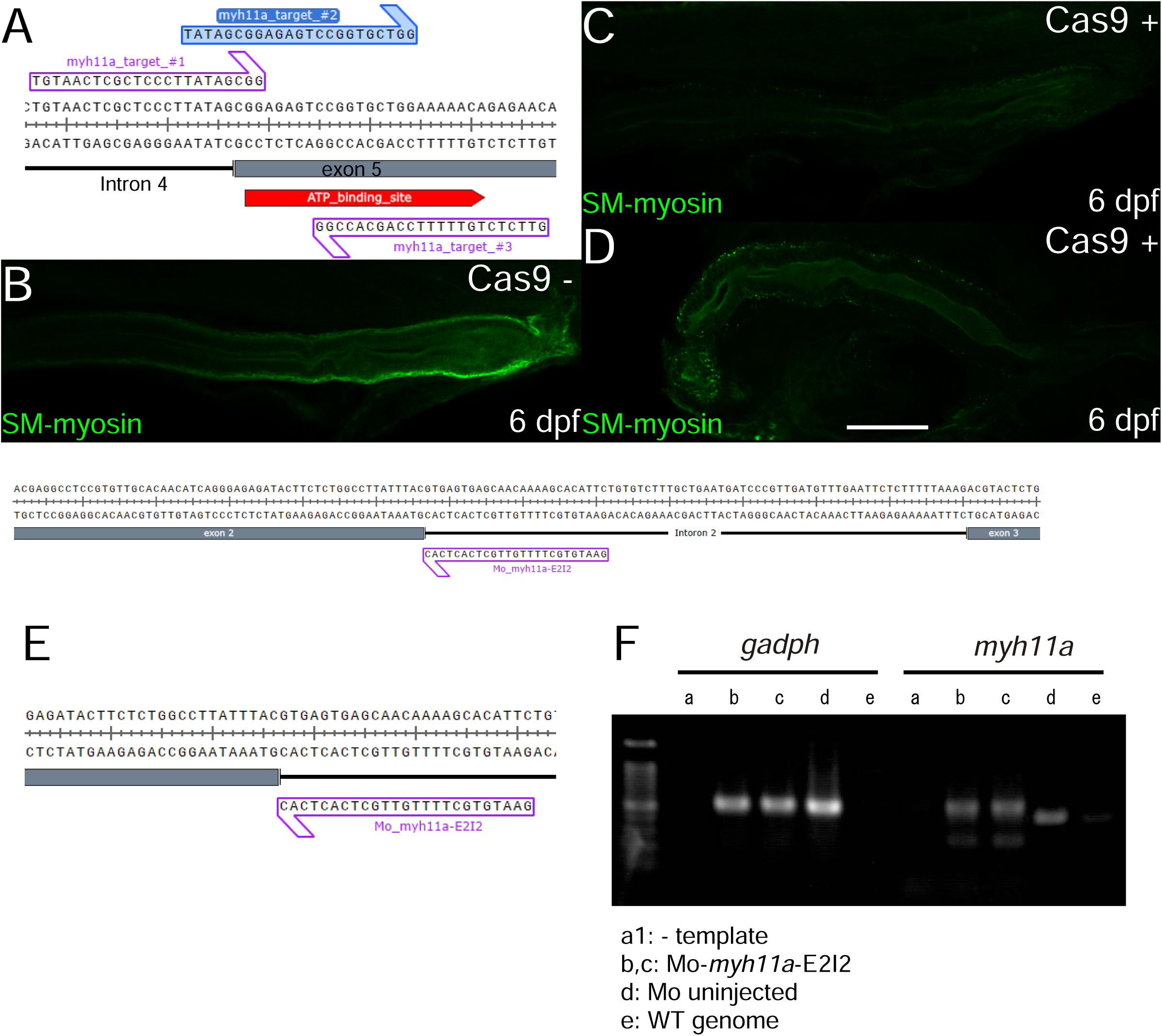
Targeting of SM-myosin *myh11a*. (A) CRISPR target sites within the *myh11a* gene. (B–D) Expression of SM-myosin in 6 dpf embryos, into which *myh11a* sgRNA was injected at 1-2 cell stage,: without Cas9 nuclease (B); with Cas9 (C and D). Scale bar: 100 µm. (E) Morpholino target site within *myh11a*. (F) RT-PCR analysis of *myh11a* morpholino-injected embryos. Lane 1: control; lanes 2 and 3: morpholino-injected; lane 4: *WT* cDNA; lane 5: *WT* genomic DNA

**Table 4.** Intestinal atresia phenotype in homozygous *myh11a* mutant embryos injected with sgRNAs with or without Cas9 nuclease.

| Cas9(+) |  |  | Cas9(-) |  |  |
| --- | --- | --- | --- | --- | --- |
| IA(+) | IA(-) | Total | IA(+) | IA(-) | Total |
| 6 | 4 | 10 | 9 | 0 | 9 |

Embryos were injected at the one-cell stage with a mixture of three sgRNAs targeting *myh11a*, either alone or in combination with Cas9 protein. The presence or absence of intestinal atresia was scored in mutant homozygous embryo. Injection of sgRNAs with Cas9 partially rescued the atresia phenotype, whereas sgRNAs injection alone had no significant effect.

**Table 5.**
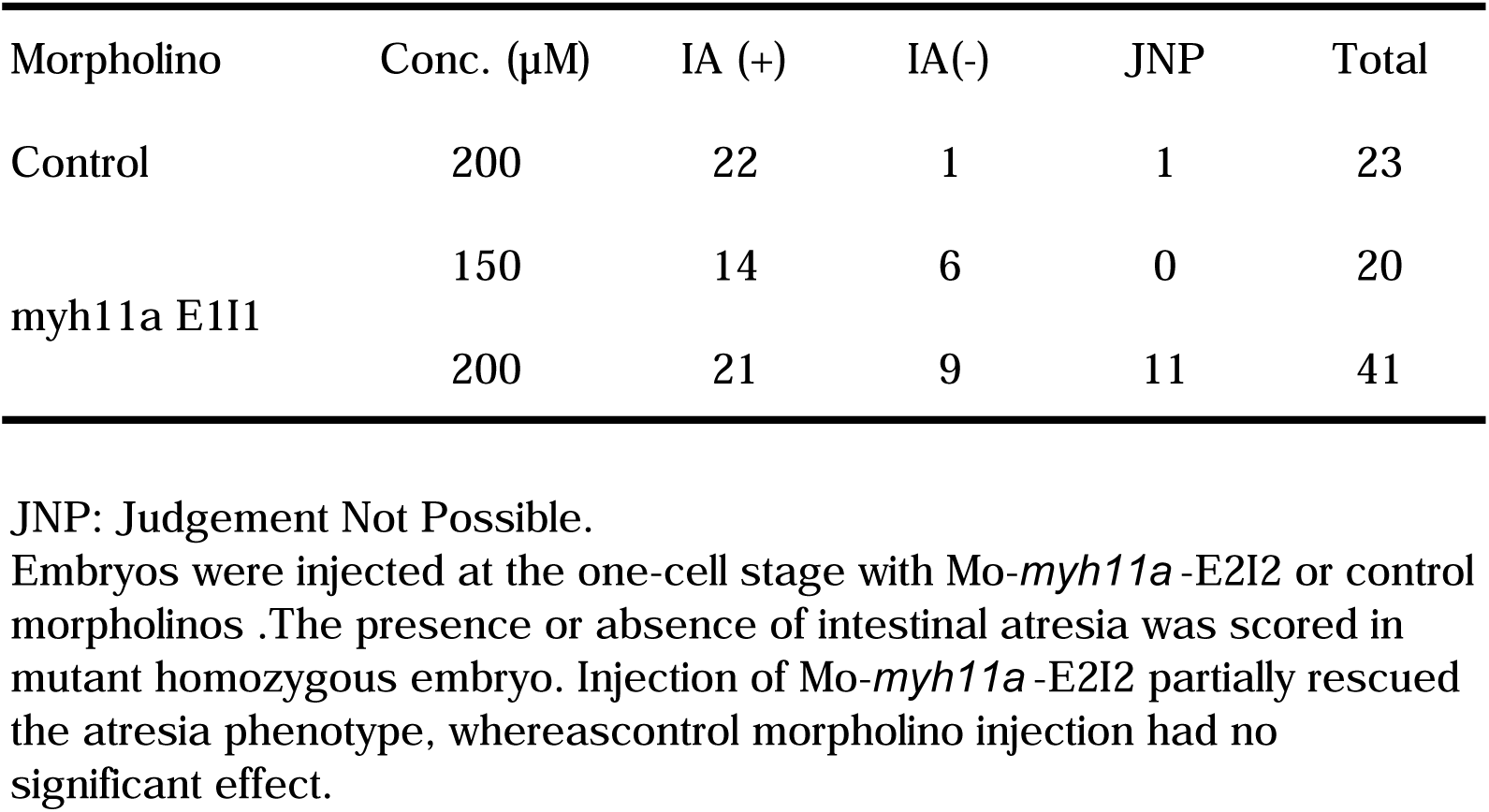
Intestinal atresia phenotype in homozygous *myh11a* mutant embryos injected with Mo-*myh11a* -E2I2 morhplino.

### Augmented Actomyosin Contraction Is Responsible for IA in *mypt1* Mutants

To further investigate whether enhanced cytoskeletal contractility contributes to the development of intestinal atresia (IA) in *mypt1* mutants, we tested whether blebbistatin—a non-muscle myosin II (NMII) inhibitor—could rescue the IA phenotype in *mypt1#4-6* mutant embryos. Embryos were treated with 50 µM blebbistatin at st. 25 for two hours (Fig. 12A). Under these conditions, no major morphological abnormalities were observed other than a pronounced curly tail phenotype (Fig. 12B, C). Remarkably, IA was not observed in all treated *mypt1* mutant embryos (Fig. 12D), strongly suggesting that excessive actomyosin contractility is a key driver of IA formation.

**Fig. 12.**
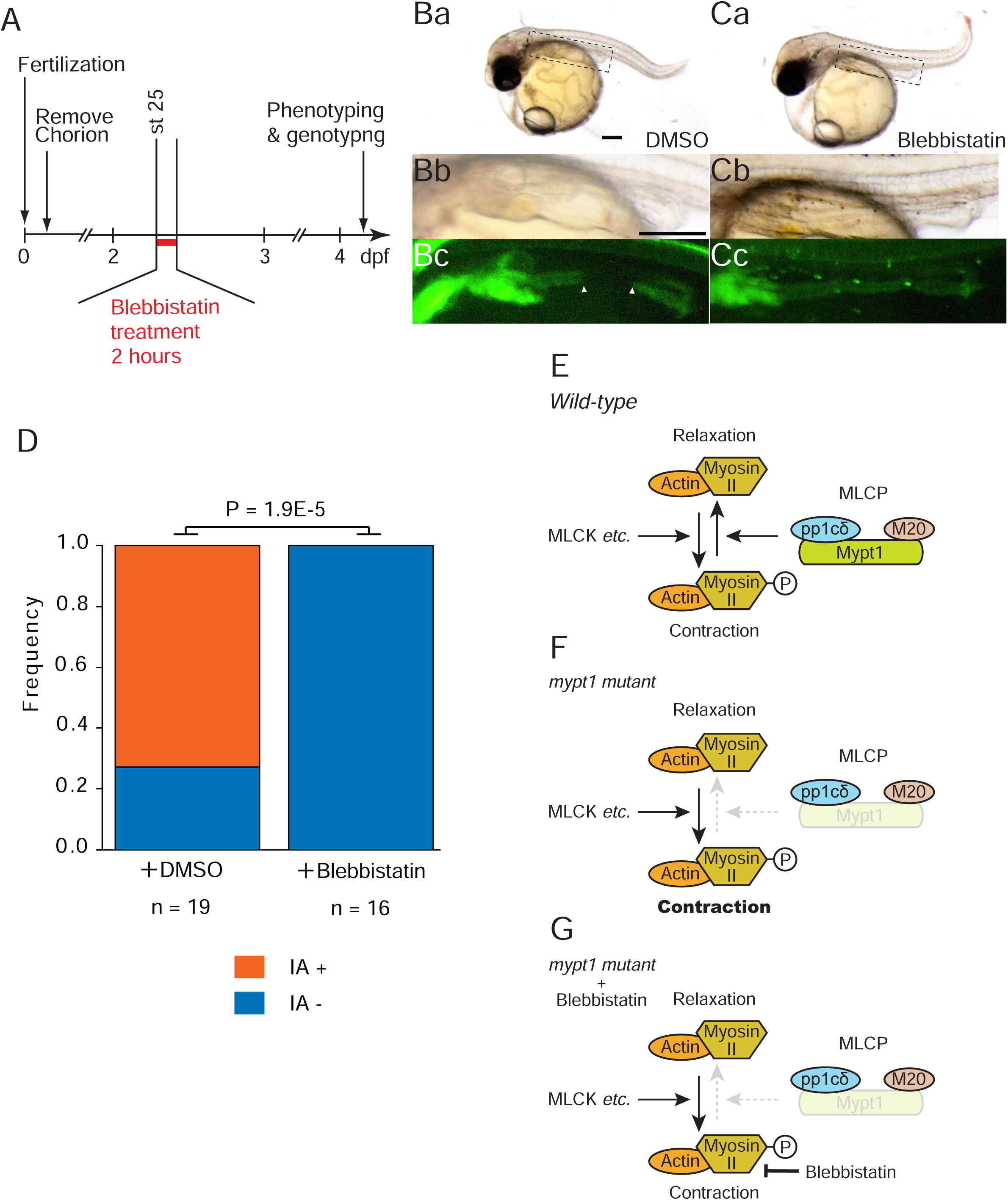
Attenuation of actomyosin activation rescues IA phenotype in *mypt1* mutants. (A) Experimental design for blebbistatin treatment. (B, C) Images at 9 dpf: control (B) and blebbistatin-treated. Arrowheads indicate IA. Scale bars: 200 µm. (Ba, Ca) Dotted boxes in (B) and (C) are enlarged in (Bb) and (Cb), respectively. (Bc, Cc) Corresponding fluorescent images of (Bb) and (Cb), respectively. (D) Quantification of rescue of the IA phenotype in *mypt1^#4-6^* embryos following blebbistatin treatment (Fisher’s exact test, *P* = 1.9 × 10^−5^). (E) Diagram of NMII regulation. *Mypt1* is a critical subunit of MLCP and is essential for its function. Loss of *Mypt1* leads to reduced MLCP activity, resulting in ectopic activation of actomyosin contraction. (F) Hypothetical molecular events following the loss of *mypt1* function. (G) Schematic representation of the molecular state under blebbistatin treatment.

## Discussion

Here, we report a novel animal model of IA. Through ENU mutagenesis of medaka, we isolated the *g1-4* mutant, in which early intestinal development proceeds normally, but loss of epithelial integrity leads to IA after the intestinal lumen has formed. Positional cloning identified a loss-of-function mutation in the *mypt1* gene. A genome-edited *mypt1* mutant allele (*mypt1^#4-6^*) recapitulated the *g1-4* phenotype, confirming that loss of Mypt1 function is responsible for IA. In *mypt1* mutants, we observed elevated levels of pMrlc and abnormal accumulation of F-actin in the developing intestine (Fig. 10). Treatment with blebbistatin, a non-muscle myosin II (NMII) inhibitor, significantly suppressed IA in *mypt1* mutants (Fig. 12), suggesting that hyperactive cytoskeletal contraction is a likely cause of epithelial disruption and IA formation. Given the limited understanding of the molecular pathophysiology of human IA and the scarcity of appropriate animal models, the medaka *mypt1* mutant provides a valuable platform for elucidating the mechanisms underlying this congenital condition.

A long-standing hypothesis proposes that IA arises from an accidental vascular injury during development (Louw, 1959; Louw and Barnard, 1955). However, in our medaka model, no overt abnormalities in blood circulation were observed in *mypt1^#4-6^*embryos. Moreover, prior studies have shown that medaka embryos establish steady and robust digestive tract blood flow by st. 30 (Fujita et al., 2006), a time point after IA formation begins. These observations suggest that IA in *mypt1* mutants does not result from vascular disruption, but instead arises from non-vascular mechanisms, such as dysregulated actomyosin contractility and impaired epithelial integrity during intestinal morphogenesis.

The developmental role of Mypt1 has been documented in some species. In *Drosophila*, *mypt1* mutants have various developmental defects, including failure of dorsal closure.(Mizuno et al., 2002; Tan et al., 2003) In these mutants, ectodermal cell sheet movement is impaired and the subcellular localization of actomyosin regulatory proteins, such as pMrlc and actin, is perturbed.(Mizuno et al., 2002) In zebrafish, *mypt1* mutants display impaired migration of bone morphogenetic protein 2a (*bmp2a*)-expressing lateral plate mesoderm (LPM) cells, causing agenesis/hypoplasia of the liver and exocrine pancreas.(Dong et al., 2019; Huang et al., 2008) Brain ventricle formation is also affected in the zebrafish mutant because of ectopically induced tension in the neuroepithelium layer.(Gutzman and Sive, 2010) In these mutants, increased pMrlc levels and aberrant localisation of NMII and F-actin are observed in the neuroepithelium. Importantly, the upregulation and mislocalisation of actomyosin regulatory components, such as pMrlc, NMII and F-actin, are likely to be common features in *mypt1* mutants across species, including medaka. Furthermore, in the context of brain ventricle development in zebrafish, blebbistatin treatment to inhibit MNII successfully rescued the *mypt1* mutant phenotype. This supports the notion that “epithelial relaxation” regulated by Mypt1 is probably critical for proper tube inflation, as occurs in brain ventricle formation. Our findings are consistent with the view that loss of *mypt1* function affects actomyosin dynamics during embryogenesis.

Despite these commonalities, notable differences exist between *mypt1* mutants in medaka and previously reported *mypt1* mutants in zebrafish. IA was not observed in zebrafish mutants, whereas neither liver nor brain ventricle abnormalities were found in medaka mutants. Although both species show F-actin accumulation in endodermal cells, the intestine of zebrafish mutants exhibits only a mildly reduced size without significant morphological changes (Huang et al., 2008). Interestingly, treatment with a BMP inhibitor phenocopies the liver-loss phenotype but does not affect intestinal size in zebrafish, suggesting that mis-localisation of *bmp2a*-expressing lateral plate mesoderm (LPM) cells is primarily responsible for the liver phenotype. In medaka, the penetrance of the IA phenotype was incomplete (Table 2). Our data indicate that differential expression of *mypt* family genes does not explain the intestinal specificity, implicating genetic background as a modifier of IA manifestation. These findings raise the possibility that genetic modifiers contribute to the variability and expressivity of the IA phenotype in medaka and may underlie interspecies differences.

Supporting this, individuals with MYPT1 mutations show highly variable clinical presentations (Contreras-Capetillo et al., 2024; Hughes et al., 2020; Picard et al., 2022), and IA has been reported in only a subset of cases. Notably, Picard et al. (2022) described a patient with IA at birth and persistent Müllerian duct syndrome (PMDS), who remained otherwise healthy until age 29. This observation is consistent with our finding that a small number of medaka *mypt1* homozygous mutants can survive to adulthood, although this is rare. During the establishment of our mutant lines, we selectively bred individuals in which IA was frequently observed, and other developmental defects were relatively rare. This selective breeding likely enriched a genetic background predisposed to IA but less susceptible to other phenotypes. Given the more than 80 established laboratory medaka strains with diverse geographic and genetic origins (Katsumura et al., 2019), it would be interesting to cross our *mypt1* mutant with other genetic backgrounds to explore the influence of genetic modifiers on IA penetrance and to reveal additional phenotypes associated with *mypt1* disruption..

Mypt1 acts as a scaffold to provide a platform for assembly of the regulatory and catalytic subunits of the MLCP complex and for the recruitment of phosphatase substrates such as NMII. Conserved domains of Mypt1 that are vital for such molecular interaction are also found in medaka Mypt1. Our original *g1-4* mutant provides insight into one such domain: C2952A mutation introduces a premature stop codon after amino acids (aa) 983, thereby deleting a large part of the Prkg interacting domain of the *WT* 1078-aa protein (Fig 1D). The LZ motif of this domain specifically mediates the interaction with Prkg1α.(Surks et al., 1999) In vascular SM cells, activation of Mlcp mediated by Prkg dephosphorylates pMrlc, which in turn induces vascular SM cell relaxation.(Surks et al., 1999) Thus, loss of the LZ motif could explain increased contraction in the *g1-4* mutant. However, it remains to be clarified whether Prkg1α or another PRKG family members regulate intestinal epithelium integrity in developing intestine.

Our experiments using CRISPR-mediated knockout and morpholino knockdown (Mo-KD) of *myh11a* suggest potential involvement of SM-myosin in IA pathogenesis. Yet, SM-myosin expression was not detected in the intestinal region where IA typically forms, suggesting that the SM layer is either absent or immature at the time of IA onset. MYH11 encodes the SM-specific myosin heavy chain, a key component of the contractile machinery required for force generation and structural stability in SM cells. However, the lack of fully developed SM cells at the timing of IA formation argues against a direct contribution of MYH11-mediated contraction of SM cells to the initial pathogenesis. In zebrafish, the *meltdown* (*mlt*) mutant— harbouring a constitutively active form of Myh11—develops cystic expansion of the posterior intestine. (Wallace et al., 2005b) This phenotype is thought to arise from Myh11-dependent stromal–epithelial signalling that disrupts epithelial architecture. A similar mechanism may operate in medaka *mypt1* mutants, where loss of Mypt1 could enhance Myh11a activity. However, several distinctions suggest divergent downstream pathways: *snail* family genes are upregulated in zebrafish *mlt* mutants but not in medaka; and whereas the *mlt* phenotype arises at 74 dpf —after SM encasement—the IA phenotype in medaka emerges by st. 28, before smooth muscle cslls are identified surrounding the region where IA develop. (Gays et al., 2017; Wallace et al., 2005a) Thus, although Myh11a dysregulation and stromal–epithelial signalling may be involved in both models, their timing and molecular mediators differ.

Interestingly, MYH11 also functions in contractile non-SM cells such as myofibroblasts, which can exert mechanical forces during development and repair (Pakshir et al., 2020). It is possible that *myh11a* functions in mesenchymal cells other than SM, contributing to mechanical stress in the developing gut and influencing epithelial morphogenesis. Although speculative, this possibility warrants further investigation. In addition, SM-specific *Mypt1* knockout (Mypt1^SMKO^) mice showed altered contractility but no apparent intestinal malformations or motility defects (He et al., 2013), further supporting the notion that canonical SM contraction is not directly responsible for IA pathogenesis. Together, these findings suggest that *myh11a* may promote IA through a SM-independent mechanism, potentially involving mechanically active mesenchymal populations such as myofibroblasts.

In addition to MYPT1 mutations, ACTB mutations have also been reported to cause IA (Fakhro et al., 2019; Saskin et al., 2017; Sibbin et al., 2021), highlighting the importance of tightly regulated actomyosin dynamics during intestinal development. Notably, mutations in ACTB and ACTG1 are known to cause Baraitser-Winter Syndrome 1 (BRWS1, OMIM #243310) in human, in which IA is an uncommon manifestation. This suggests that while disruption of actomyosin dynamics contributes to IA pathogenesis, it is not solely sufficient to induce the condition. Supporting this, pharmacological inhibition of NM II using blebbistatin completely rescued the IA phenotype in *mypt1* mutants, whereas treatment with calyculin A treatment—used to inhibit PP1 and thus supress dephosphorylation of NMII— failed to phenocopy the IA phenotype. This discrepancy implies that Mypt1 may exert additional, non-canonical functions beyond its regulatory role in MLCP (Kiss et al., 2019), or that inhibition of PP1 alone is is insufficient to increase the amount of phosphorylated posho-MII and contractility ofin intestinal epithelial cells. Identification of genetic modifiers that influence IA development in the *mypt1* mutant background may provide further mechanistic insight.

Our data show that ectopic accumulation of pMrlc and F-actin, likely indicative of actomyosin hypercontractility, occurs in a non-uniform pattern along the intestine in *mypt1* mutants. The cause of this spatial variability remains unknown. Moreover, we have not yet validated the molecular mechanism that underlies how hypercontractility in the intestinal epithelium disintegrates the epithelial layer structure. Further study is required to assess completely the mechanics underpinning embryonic epithelial integrity.

### Experimental Procedures

#### Ethics statement and Fish Strains

All experiments were approved by the animal experimentation committee of Kyoto Prefectural University of Medicine (KPUM), licence numbers 27-123 and 2019-154. Fish were kept at 26 °C on a 14 hours light / 10 hours dark cycle in a constant re-circulating system of the in-house KPUM facility. The d-rR medaka strain original *mypt1* mutant *g1-4* and hatching enzyme were obtained from NBRP (National Institute for Basic Biology, Okazaki, Japan). Embryos were staged according to Iwamatsu’s staging system.(Iwamatsu, 2004)

#### Knock-in transgenic (KIT) medaka

To visualise the intestinal epithelium, we generated a KIT medaka line that expresses membrane-bound GFP (memGFP) under the control of the endogenous *foxa2* promotor (Fig. 3E), *Tg*[*foxa2:memGFP*]. *foxa2* is a well-known marker for the endoderm and its derived organs.(Kobayashi et al., 2006) The targeting vector was constructed as described.(Ansai and Kinoshita, 2017; Okuyama et al., 2013) This construct was integrated into the genome as previously described.(Murakami et al., 2017) The sgRNA for the genomic target site was designed as previously described (Kimura et al., 2014).

#### Detection of Apoptosis by Acridine Orange and TUNEL Assay

To detect apoptotic cells, acridine orange (AO) staining and TUNEL assays were performed. For AO staining, a stock solution of AO (0.05 mg/mL; Tokyo Chemical Industry, Tokyo, Japan) was prepared in distilled water and stored at 4°C in the dark. Prior to use, the stock solution was diluted to 17 μg/mL in hatching buffer (HB). Embryos at st. 27 were dechorionated and incubated in the AO solution for 30 minutes, then washed three times in fresh HB for 5 minutes each to remove excess dye.

Fluorescent images were acquired using a DP70 digital camera mounted on an SZX16 stereomicroscope (Olympus, Tokyo, Japan). TUNEL assays were performed using the *In Situ Apoptosis Detection Kit* (Takara) according to the manufacturer’s instructions. To visualise the intestinal epithelial membrane, embryos carrying the *Tg[foxa2:memGFP]* transgene were used. GFP signals were detected using a rabbit anti-GFP antibody (Invitrogen, A6455), followed by a goat anti-rabbit IgG antibody conjugated with Alexa Fluor 633 (Thermo Fisher Scientific).

#### Whole mount *in situ* hybridization

A *foxa2* probe was designed and whole mount *in situ* hybridisation was performed as described previously.(Kobayashi et al., 2006) (Takashima et al., 2007) Template DNAs for *snai1a* (EST clone number, olea55k06), *snai1b* (olea23g07) and *snai2* (olsp48a08) were obtained from the National BioResorce Project (NBRP) medaka (https://shigen.nig.ac.jp/medaka/).

#### Histology

Embryos fixed with 4% paraformaldehyde (PFA) were embedded in paraffin or resin (Technovit 7100; Heraeus, Werheim Germany) and 6 µm tissue sections were prepared. Paraffin sections were stained with haematoxylin and eosin, while plastic sections were stained with haematoxylin only, followed by analysis using a BX51 microscope (Olympus, Tokyo, Japan).

#### Immunohistology

For staining SM myosin, laminin and cytokeratin, 80% methanol/20% DMSO (Dent’s solution) was used for fixation. In other cases, embryos were fixed in 4% PFA in 1.5× PBS containing 0.1% Tween 20. If Dent’s fixation was used, embryos were de-chorionated with medaka hatching enzyme before fixation. Then, non-specific antibody binding sites were blocked with 1% dimethyl sulfoxide, 2% bovine serum albumin, 10% normal goat serum, 0.1% Triton X-100 in 1× PBS. Embryos fixed in PFA were permeabilised by incubation in 1× PBS containing 2% Triton X-100 for at least 2 hours before blocking of non-specific binding sites. The following commercially available antibodies were used: GFP (1/2,000; ab13970; Abcam), cytokeratin (1/100; clone AE1/AE3, ab27988; Abcam), SM myosin (MYH11, 1/50; BT562; Biomedical Technologies), laminin (1/100; L9393; Sigma), phospho-myosin light chain 2 (Ser19) (1:20; #3671; Cell Signaling Technology), phospho-Histone H3 (Ser10) (1/500; #06-570, Millipore), and PKCζ (1/800; ab5964; Abcam,). Alexa Fluor-633-Phalloidin (1/20; A22284; Thermo Fisher Scientific) was used to mark F-actin. Detection of primary antibodies was performed using Alexa Fluor-488, -555 goat anti-rabbit, chicken or mouse IgG (1/400; Invitrogen). Images were acquired with an Olympus FV1200 confocal microscope or Zeiss LSM900 and yz-planes were reconstructed using FIJI.(Schindelin et al., 2012) Abnormal accumulation of pMrlc or F-actin was judged based on: (a) obviously higher immunofluorescence signal in the middle part of the intestine compared with other areas, and (b) obviously higher immunofluorescence signal at apical and also basal areas of the intestinal epithelium where strong signal was never observed in the *WT*. Representative single-plane confocal images (not z-projections) are shown in the figures.

#### CRISPR/Cas9-Mediated Mutagenesis of *mypt1*

The sgRNA targeting exon 1 of *mypt1* was designed using the web tool “Search for CRISPR target site with micro-homology sequences” (http://viewer.shigen.info/cgi-bin/crispr/crispr.cgi), with the parameter “micro-homology sequences” set to 4 bases. sgRNA synthesis was performed as previously described.(Ansai and Kinoshita, 2014) The target sequence is shown in Fig. 2A. A mixture containing 150 ng/µL sgRNA and 200 ng/µL Cas9 mRNA was microinjected into 1–2 cell stage embryos. Injected embryos were raised to adulthood, and eight germline-transmitting F0 founders were identified by genotyping pooled F1 offspring. One of these founders, #4-6, was outcrossed to *WT* fish, and the resulting F1 progeny were genotyped. F1 fish harbouring the mutation (Fig. 1, 2) were used for subsequent analyses.

#### CRISPR/Cas9 knockout of *myh11a*

To knock out *myh11a*, three overlapping sgRNAs targeting exon 5—which encodes part of the ATP-binding domain—were designed using the CRISPR-Cas9 guide RNA design checker (Integrated DNA Technologies, IDT). Off-target effects were evaluated via BLAST search against the medaka genome using the Ensembl genome browser. The overlapping design of sgRNAs was intended to increase mutagenesis efficiency in the targeted region (Fig. 11A). Each sgRNA was prepared following the manufacturer’s protocol (IDT). A mixture containing 1.0 µg/µL Alt-R™ S.p. Cas9 Nuclease V3 (IDT) and 5.0 µM of each sgRNA, diluted in RNase-free water was injected into 1–2 cell stage medaka embryos.

#### Morpholino knockdown of myh11a

Antisense Morpholino (Gene Tools) targeting the splice donor site of exon 2of medaka *myh11a* was used for knockdown (Fig. 11E). The Standard Control Oligo supplied by Gene Tools was used as a negative control.

#### Genotyping

*WT* and mutant *mypt1* alleles were distinguished using a polymerase chain reaction-restriction fragment length polymorphism (PCR-RFLP) method. Genomic DNA from lysed tissue was subjected to PCR using Quick Taq® HS DyeMix Taq polymerase (TOYOBO), with an initial denaturation at 94°C for 2 min, followed by 45 cycles of 94°C denaturation for 30 s, annealing at 55°C for 30 s, and extension at 68°C for 50 s,. The primers used are shown in Table 6. PCR products were digested with the restriction enzyme *Hae*III (TOYOBO), and analysed by electrophoresis in 3% agarose gels (02468-95, Nacalai) in 1× TAE buffer.

**Table 6.**
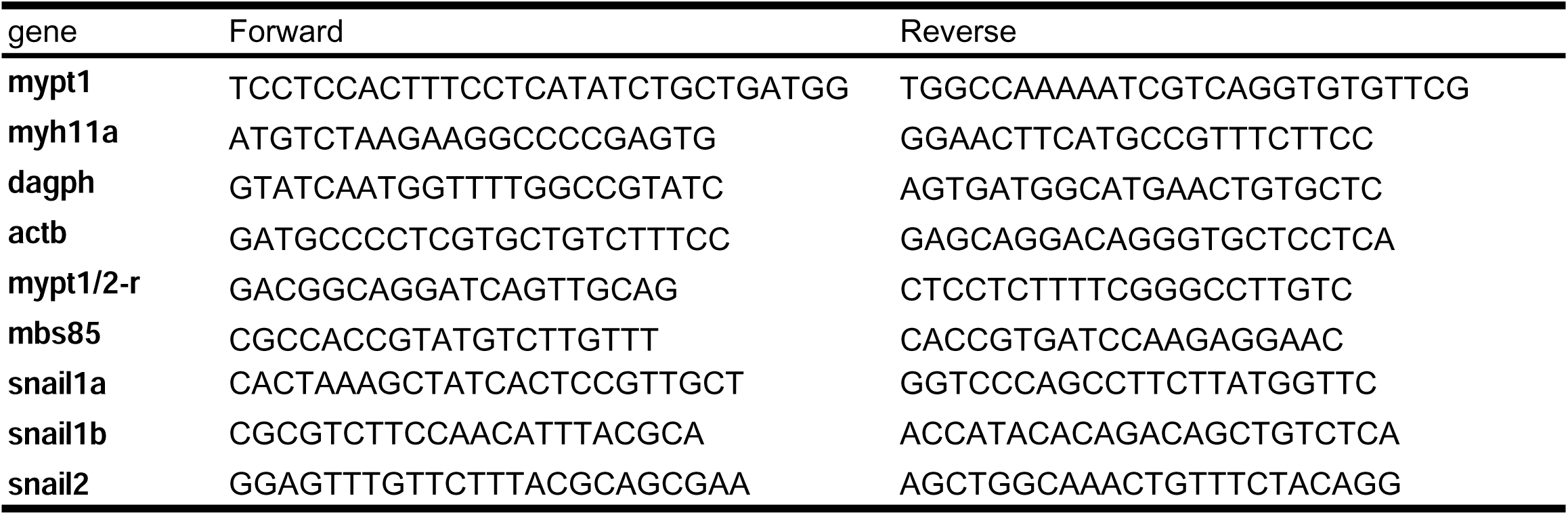
Primers.

#### Chemical Inhibition of Actomyosin Contractility

To examine the effect of NMII inhibition on IA development, st. 25 embryos were treated with either 0.1% DMSO (control) or 50 µM (-)-blebbistatin (021-17041, FUJIFILM), dissolved in 0.1% DMSO. After a 2-hour incubation, embryos were thoroughly washed and raised until 4 dpf for phenotypic analysis.. Furthermore, embryos were raised to 9 dpf for extended observation, imaging and genotyping. To assess the effect of increased myosin phosphatase inhibition, *WT* embryos were treated from st. 23 to st. 32 with either 0.1% DMSO (control) or 0.1 µM Calyculin A (101932-71-2, FUJIFILM), also dissolved in 0.1% DMSO. Phenotypic evaluation was performed after the treatment, and embryos were further raised to 9 dpf for evaluation.

#### Phylogenetic analysis

Multiple sequence alignments were performed using CLUSTALW with default parameters, followed by phylogenetic tree construction using the PhyML-bootstrap method (Kyoto University Bioinformatics Center, https://www.genome.jp/tools-bin/clustalw). The protein sequences used for the alignment are listed in Table 3.

#### RT-PCR

Total RNA was extracted using TRIzol reagent (Invitrogen) according to the manufacturer’s protocol. For small tissue samples, RNA was purified using the Direct-zol RNA Kit (Zymo Research). Reverse transcription and PCR were carried out using superscript III (Invtrogen) and Quick Taq® HS DyeMix (TOYOBO) , respectively, following the manufacturer’s instructions.

#### Statistical analysis

Statistical analyses were performed using EZR(Kanda, 2013)

## Acknowledgements

The medaka National Bioresource Project (NBRP: https://shigen.nig.ac.jp/medaka/), governed by the Ministry of Education, Culture, Sports, Science and Technology, Japan, provided the hatching enzyme, EST clones, d-rR and original mutant *g1-4* medaka strains used in this study.

## Author Contributions

DK designed the research; DK, AK, TK, KM, SA, MK, and KN examined the mutant phenotype; DK, TK, HY, ST, TK, TK, TN and TJ performed mutant screening; YI, KA, YS and HT organized mutant screening; DK, KM, YN, MS, SS, SI, TY, HT and KY analysed data; and DK and KY wrote the paper.

